# Structural Definition of Neutralizing Epitopes within and around the HKU1 Receptor-binding Site Reveals Cell Context–Dependent Neutralization

**DOI:** 10.1101/2025.10.29.685445

**Authors:** Jeswin Joseph, Lingshu Wang, Sheena Vasquez, Chia-Ming Su, Daniel Wrapp, Osnat Rosen, Olubukola M. Abiona, Christian K.O. Dzuvor, Robert N. Kirchdoerfer, Sydney P. Moak, Timothy P. Sheahan, Catherine Hammond, Wei Shi, Wing-Pui Kong, Yi Zhang, Michael R. Eso, Ariane J. Brown, Tanushree Dangi, Pablo Penaloza-MacMaster, Andrew B. Ward, Ralph Baric, Jason S. McLellan, Theodore C. Pierson, John Mascola, Barney S. Graham, Hadi M. Yassine, Mohsan Saeed, Christopher O. Barnes, Kizzmekia S. Corbett-Helaire

## Abstract

Endemic human coronaviruses (HCoVs), such as HCoV-HKU1, account for ∼30% of common colds each year and can cause serious upper and lower respiratory infections, yet no licensed vaccines or therapeutic antibodies target HCoVs. Despite being endemic to the human population, little is known about the antigenic landscape of HCoV-HKU1. Here, we characterized key interactions between the HCoV-HKU1 spike (S) protein and monoclonal antibodies (mAbs) isolated from convalescent HCoV-HKU-1-positive peripheral blood mononuclear cells collected prior to the COVID-19 pandemic. We isolated 14 mAbs that bound distinct regions of S, including the receptor-binding domain (RBD), N-terminal domain, and S2 subunit. Structural and functional studies revealed three groups of RBD-specific mAbs targeting diverse footprints within and around the TMPRSS2 receptor-binding site: (1) H501-022, which recognizes the TMPRSS2-binding site and thereby blocks receptor engagement; (2) H501-008, which binds a distinct epitope outside the TMPRSS2-binding site that is shared with HCoV-OC43; and (3) H501-018, which recognizes both “up” and “down” RBD conformations at a distinct, non-overlapping epitope outside the TMPRSS2-binding site. H501-008 weakly neutralized live HCoV-OC43 *in vitro* and protected mice against lethal HCoV-OC43 challenge. Notably, the three RBD-specific mAbs potently neutralized HCoV-HKU1 pseudovirus in TMPRSS2-overexpressing cell lines, but exhibited limited neutralizing activity against authentic HCoV-HKU1 infection in primary human airway epithelial cells. Together, these findings structurally define the antigenic landscape within and around the HKU1 receptor-binding site and demonstrate that receptor-binding site recognition alone is insufficient to predict physiologically relevant antibody-mediated neutralization.

## Introduction

Coronavirus disease 2019 (COVID-19), caused by severe acute respiratory syndrome coronavirus 2 (SARS-CoV-2), has claimed more than seven million lives globally, emphasizing the persistent threat posed by coronaviruses (CoVs)^1^. However, four endemic human coronaviruses (HCoVs), HCoV-HKU1, HCoV-OC43, HCoV-229E, and HCoV-NL63, are responsible for up to 30% of seasonal common colds^2^. HCoV infection typically manifests as asymptomatic or mild upper respiratory disease^3^, but can also result in hospitalization and death in older adults, individuals with co-morbidities, or immunocompromised populations^3–6^.

Despite widespread circulation, there are currently no licensed vaccines or therapeutics for HCoVs, although the concept of incorporating HCoV antigens into supra-seasonal or universal CoV vaccines has been explored in pre-clinical studies by our group and others^7–9^. Unfortunately, such vaccine development is hindered by an incomplete understanding of the antigenic landscape of the major HCoV surface protein, spike (S). For the S proteins of Middle East respiratory syndrome coronavirus (MERS-CoV), SARS-CoV, and SARS-CoV-2, the receptor-binding domain (RBD) is the principal target of potently neutralizing antibodies and is thus the preferred CoV vaccine antigen^10^. RBDs are located within the C-terminal domain of the S1 head region and play a critical role in viral entry^11,12^, whereas the N-terminal domain (NTD) typically mediates interactions with accessory host-cell attachment factors^12,13^.

Until recently, *O*-acetylated sialic acids were recognized as attachment factors for beta-HCoVs^14^, but functional entry receptors remained unknown. A series of recent studies delineated the two-step receptor engagement process used by HCoV-HKU1 S^15,16^. First, the HCoV-HKU1 NTD binds cellular sialic acid moieties, which triggers RBD exposure^15^. Next, the HCoV-HKU1 RBD binds the functional entry receptor transmembrane protease serine 2 (TMPRSS2)^16^. This glycoprotein interaction, followed by conformational changes in the RBD, enables viral-host membrane fusion and subsequent HCoV-HKU1 infection. Structural studies of HCoV-HKU1 S revealed the occluded nature of the RBD^12^, compared with epidemic CoVs, which exhibit transient open and closed RBD conformations^13,17^. RBD occlusion may represent an immune-evasion mechanism adopted by HCoV-HKU1 that limits antibody access to this functionally critical region. Similar immune-evasion mechanisms have been reported in human immunodeficiency virus (HIV), in which primary receptor engagement induces conformational changes that facilitate interaction with a secondary receptor^18^.

Functional characterization of diverse monoclonal antibodies (mAbs) following HCoV-HKU1 infection is one approach to map the antigenic sites associated with both the open and closed states of S. Extensive global efforts have led to isolation and structural characterization of hundreds of potently neutralizing mAbs targeting diverse S epitopes of epidemic CoVs, particularly within the RBD and NTD^19,20^. These discoveries established the RBD as the principal target of potent neutralizing antibodies, revealed key mechanisms of viral neutralization, identified broadly cross-reactive antibody lineages, and directly informed the design of antibody-based therapeutics and next-generation vaccines. In contrast, systematic mAb discovery efforts for endemic HCoVs such as HCoV-HKU1 remain limited. To date, only one type-specific HCoV-HKU1 RBD-specific mAb (R7)^21^ has been structurally characterized, and it was identified through a phage-display platform based on a human naive single-chain variable fragment library. Therefore, significant gaps remain in our understanding of the antibody response to HCoV-HKU1 infection and the antigenic landscape of the intact spike protein, knowledge that could be leveraged to design vaccines with broader CoV protection.

Here, we used human HCoV-HKU1-positive peripheral blood mononuclear cells (PBMCs) acquired prior to the COVID-19 pandemic and a rational B-cell sorting strategy to isolate a panel of mAbs. The resulting 14 mAbs targeted distinct regions of the S protein, including the RBD (N = 11), NTD (N = 2), and S2 stem domain (N = 1). Anti-HKU1 RBD-specific mAbs were categorized into three major competition groups and assessed for pseudovirus neutralization activity in cell lines, suggesting multiple mechanisms of action. Structures of neutralizing anti-HKU1 RBD-specific mAbs from each competition group bound to prefusion-stabilized HCoV-HKU1 S (S-2P)^22^ revealed discrete sites of vulnerability within and around the TMPRSS2 receptor-binding site and provide a molecular framework for understanding HCoV neutralization mechanisms.

Importantly, the anti-HKU1 RBD mAbs did not effectively neutralize authentic viral isolates in primary human airway epithelial cells, suggesting that physiologically relevant neutralization mechanisms differ from mechanisms of pseudovirus neutralization in TMPRSS2-overexpressing cell lines. Together, these findings report the isolation and characterization of the first type-specific HCoV-HKU1 mAbs from a convalescent donor, define key antigenic features of HCoV-HKU1 S, advances understanding of similarities and differences between antibody neutralization of epidemic and endemic HCoVs, and challenges the understanding that RBD-specific mAbs, especially those that block receptor engagement, are typically potent neutralizers. These findings will improve understanding of endemic CoV immune imprinting and facilitate assessment of universal vaccine-elicited immune responses towards broadening the immunological reach of future vaccine designs.

## Results

### HCoV-HKU1-positive convalescent PBMCs yield mAbs that bind various S domains

Human donor A from the VRC-200 clinical study presented with acute HCoV-HKU1 infection in 2014 (6 years prior to the COVID-19 pandemic); sera and PBMCs were collected during early convalescence, 41 days after diagnosis. Serum reacted with both HCoV-HKU1 and HCoV-OC43 S proteins, but not SARS-CoV or MERS-CoV, likely reflecting prior exposure to endemic HCoVs during the donor’s lifetime^23^ **(Figure 1A)**. We therefore used HCoV-HKU1 S-2P probes to isolate antigen-specific B cells, and probe-positive cells were subsequently sequenced to identify antibody heavy- and light-chain sequences for 14 mAbs (H501-003, -007, -008, -009, - 010, -012, -013, -014, -015, -016, -018, -020, -022, and -101). **(Figure 1B and Table 1)**.

**Figure 1.**
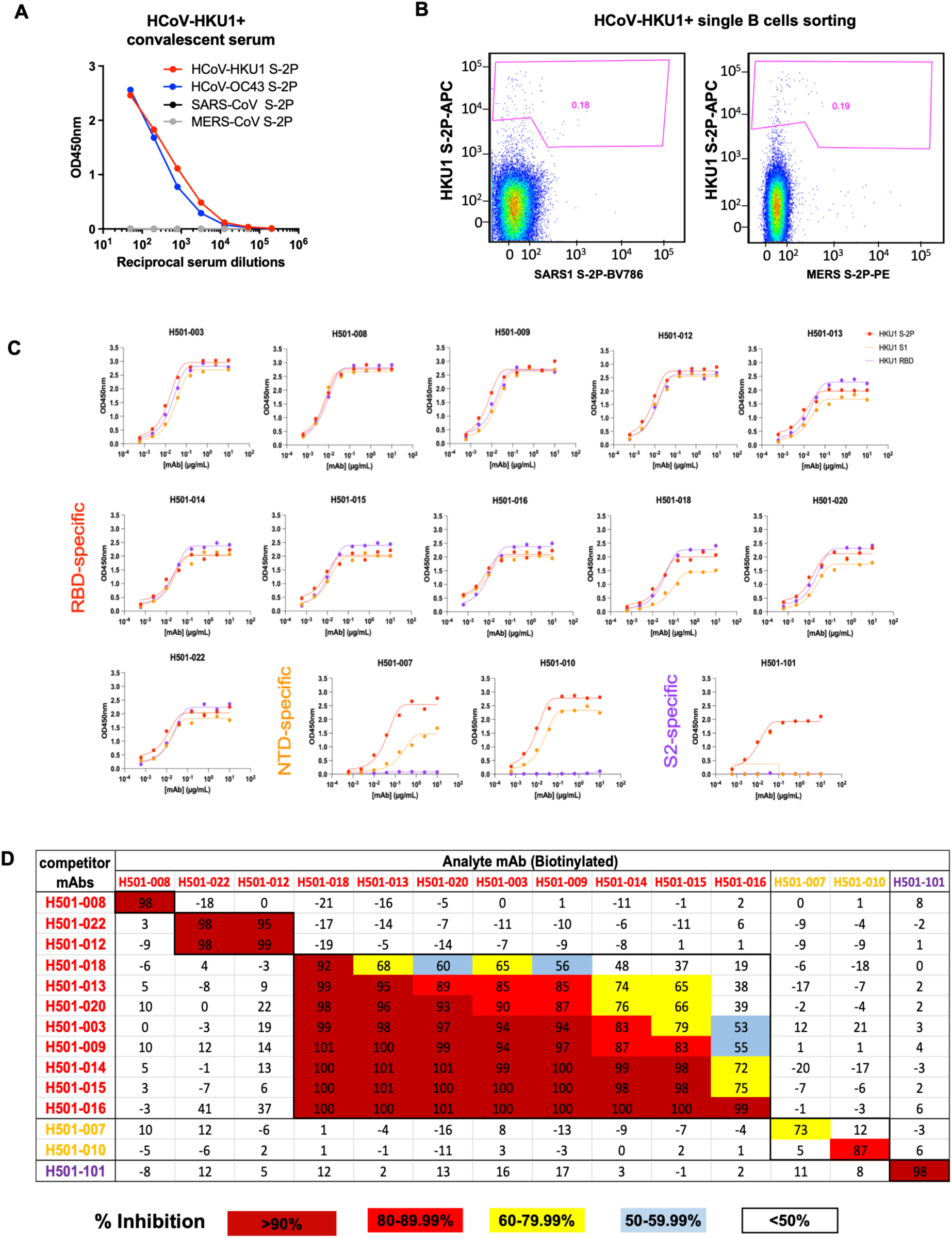
HCoV-HKU1 S-specific mAb isolation strategy and mAb competition analysis. **(A)** Human serum collected 41 days after a confirmed HCoV-HKU1 infection was assessed for binding to HCoV-HKU1 (red), HCoV-OC43 (blue), SARS-CoV (black), and MERS-CoV (gray) S-2P proteins by ELISA. SARS-CoV and MERS-CoV data points are overlaid on the x-axis. **(B)** PBMCs collected 41 days after a confirmed HCoV-HKU1 infection were stained with a viability marker (VIVID), Abs to exclude T cells and monocytes (CD3/CD4/CD8/CD14), Abs to label B cells (CD20/IgG), and S-2P probes for HCoV-HKU1, SARS-CoV, and MERS-CoV. IgG^+^ HCoV-HKU1 S-2P single B cells were sorted and subjected to PCR amplification and cloning of V_H_/V_L_ genes. A panel of mAbs was generated and selected for further characterization based on binding specificity and neutralization potency (Table 1). **(C)** Binding of HCoV-HKU1 mAbs to HCoV-HKU1 S-2P (red), S1 (orange), and S2 (purple) by ELISA. mAbs were determined to be RBD-specific if they bound all three proteins. mAbs were determined to be NTD-specific if they bound S-2P and S1. mAbs were determined to be S2-specific if they bound only S-2P. **(D)** HCoV-HKU1 RBD-(red), NTD-(orange), and S2-specific (purple) mAbs were biotinylated and assessed for competition binding to HCoV-HKU1 S-2P bound by mAbs listed in the leftmost column. Analyses were completed by ELISA. Percent inhibition of analyte (biotinylated) mAb binding by competitor (unlabeled) mAb is indicated by color. Dark red = >90% competition, red = 80–99.99% competition; yellow = 60–79.99% competition; light blue = 50–59.99% competition; uncolored = <50% competition.

**Table 1.** HCoV-HKU1 mAb binding specificity, affinity, and neutralization potency. ^1^Binding of mAbs was assessed by ELISA to HCoV-HKU1 S-2P, S1, and RBD proteins. EC50 was calculated. “-” = no curve generated. ^2^ Affinity of mAbs bound to proteins, per Extended Data Fig. 3, was measured using biolayer interferometry and K_D_, Ka and Kd were calculated. ^3^Neutralization activity was measured using a HCoV-HKU1 neutralization assay. IC values represent geometric mean titers (GMT) from two independent experiments. ^4^mAbs were determined to be RBD specific if they bound to all three proteins. ^5^mAbs were determined to be NTD specific if they bound to S-2P and S1. ^6^mAbs were determined to be S2-specific if they only bound S-2P.

| Specificity | Name | ELISA EC<sub>50</sub> (μg/mL)<sup>1</sup> |  |  | Octet<sup>2</sup> |  |  | Pseudovirus Neutralization(μg/mL)<sup>3</sup> |  |
| --- | --- | --- | --- | --- | --- | --- | --- | --- | --- |
|  |  | S-2P | S1 | RBD | K<sub>D</sub> (M) | k<sub>a</sub> (M<sup>-1</sup>s<sup>-1</sup>) | k<sub>d</sub> (s<sup>-1</sup>) | IC50 | IC80 |
| RBD<sup>4</sup> | H501-008 | 0.0055 | 0.0049 | 0.0068 | 9.37E-10 | 3.36E+05 | 3.15E-04 | 0.05 | 0.12 |
|  | H501-022 | 0.0064 | 0.0144 | 0.016 | 2.01E-09 | 2.76E+05 | 5.55E-04 | 0.01 | 0.04 |
|  | H501-012 | 0.0059 | 0.0079 | 0.0097 | 9.69E-10 | 3.12E+05 | 3.02E-04 | 0.02 | 0.06 |
|  | H501-018 | 0.0191 | 0.0574 | 0.0306 | 3.70E-10 | 2.83E+05 | 1.05E-04 | 0.28 | 0.74 |
|  | H501-013 | 0.0077 | 0.0144 | 0.0155 | 9.15E-10 | 2.28E+05 | 2.09E-04 | 3.77 | >10 |
|  | H501-020 | 0.0094 | 0.0193 | 0.0172 | <1.0E-12 | 3.33E+05 | <1.0E-07 | 0.32 | 0.85 |
|  | H501-003 | 0.0122 | 0.0277 | 0.0176 | 5.08E-10 | 2.19E+05 | 1.11E-04 | 0.57 | 1.59 |
|  | H501-009 | 0.0041 | 0.0134 | 0.0095 | 5.44E-09 | 7.03E+04 | 3.83E-04 | 4.91 | >10 |
|  | H501-014 | 0.0084 | 0.0177 | 0.0188 | 2.65E-10 | 1.92E+05 | 5.10E-05 | 1.66 | 3.75 |
|  | H501-015 | 0.0041 | 0.0094 | 0.0098 | 5.68E-10 | 2.06E+05 | 1.17E-04 | 1.32 | 2.43 |
|  | H501-016 | 0.0054 | 0.0078 | 0.0079 | 1.76E-10 | 1.85E+05 | 3.26E-05 | 2.54 | 6.09 |
| NTD<sup>5</sup> | H501-007 | 0.0327 | 0.3471 | - | 2.15E-07 | 2.34E+04 | 5.04E-03 | >10 | >10 |
|  | H501-010 | 0.0076 | 0.0177 | - | 5.88E-09 | 4.78E+05 | 2.81E-03 | >10 | >10 |
| S2<sup>6</sup> | H501-101 | 0.006 | - | - | <1.00E-12 | 1.85E+05 | <1.0E-07 | >10 | >10 |

First, to determine S-region binding specificity, we assessed the ability of the mAbs to bind HCoV-HKU1 S-2P^12^, S1, and RBD proteins by ELISA. Eleven mAbs (H501-003, -008, -009, -012, -013, -014, -015, -016, -018, -020, and -022) specifically bound HCoV-HKU1 RBD. Two mAbs (H501-007 and -010) bound HCoV-HKU1 S-2P and S1, but not RBD, and were thus classified as NTD-specific. One mAb (H501-101) bound HCoV-HKU1 S-2P, but not S1 or RBD, and was therefore classified as S2-specific **(Table 1 and Figure 1C)**. With the exception of H501-003, - 007, and -018, the mAbs bound HCoV-HKU1 S-2P with half-maximal effective concentration (EC_50_) values of less than 0.01 µg/mL **(Table 1 and Figure 1C)**. Similarly, binding affinities determined by biolayer interferometry (BLI) revealed high-affinity interactions between the mAbs and their corresponding HCoV-HKU1 S domains, with H501-020 and -101 exhibiting the highest affinities for HCoV-HKU1 RBD and S-2P, respectively **(Table 1 and Figure S1)**. In general, RBD mAbs exhibited higher affinities than NTD mAbs **(Table 1)**.

We next assessed whether the HCoV-HKU1 mAbs recognized HCoV-HKU1 genotype A. NTD mAb H501-007, RBD mAbs H501-012 and H501-022, and S2 mAb H501-101 bound HCoV-HKU1-A S protein (**Figure S2**), whereas the remaining mAbs recognized only genotype C. These findings demonstrate that most of the isolated mAbs are genotype-specific, underscoring fine antigenic differences among HCoV-HKU1 genotypes that shape the strain specificity of the antibody response.

Next, we evaluated competitive binding of biotinylated anti-HKU1 mAbs to HCoV-HKU1 S-2P using ELISA **(Figure 1D)** and BLI **(Figure S3)**. RBD mAbs H501-012 and -022 showed competitive binding to HCoV-HKU1 S-2P, consistent with their close clonal relationship (i.e., sharing the same heavy- and light-chain germline genes, identical CDRH3 and CDRL3 lengths, and >90% identity across both regions) **(Figure 1D, Figure S3, and Table S1)**. H501-008 exhibited a unique competition profile and did not compete with other RBD mAbs, consistent with recognition of a distinct epitope. H501-018 showed limited competition with H501-008 and H501-022 but competed with several other RBD-specific mAbs, defining a broader competition group represented by H501-018. NTD mAbs H501-007 and -010 competed with each other but not with RBD or S2 mAbs. Likewise, the S2-specific mAb H501-101 showed no competition with either RBD or NTD mAbs **(Figure 1D and Figure S3)**. Based on these competition profiles, the RBD-specific mAbs segregated into three distinct groups represented by H501-008, H501-018, and H501-022. H501-018 was selected as a representative mAb from the second competition group for further structural and functional characterization due to its modestly higher pseudo-neutralization titers compared to competing mAbs, as detailed below **(Table 1)**. Consistent with previous studies^24,25^, our data suggest that CoV infection elicits mAbs targeting diverse S regions with varying affinities.

### Three RBD-specific mAbs potently neutralize HCoV-HKU1 pseudoviruses in monolayer cells

Next, we evaluated the neutralization potential of our mAb panel using HCoV-HKU1 pseudoviruses (genotype C, first identified in 2006)^26^. HEK293T cells expressing human TMPRSS2 were infected with HCoV-HKU1 pseudoviruses complexed with serially diluted mAbs. RBD mAbs showed a range of neutralizing activities, with H501-008, -012, and -022 exhibiting the greatest potency, with half-maximal inhibitory concentration (IC_50_) values of 0.05, 0.02, and 0.01 µg/mL, respectively. The remaining RBD mAbs exhibited little to no neutralization capacity against HCoV-HKU1, with IC_50_ values ranging from 0.28–4.91 µg/mL. In contrast, none of the NTD or S2 mAbs effectively neutralized HCoV-HKU1 pseudoviruses (IC_50_ > 10 µg/mL) **(Table 1)**. Although there are several notable exceptions^27^, CoV NTD and S2 mAbs are generally moderately neutralizing or non-neutralizing^28,29^, consistent with our observations. Importantly, these pseudovirus neutralization data demonstrate that HCoV-HKU1 RBD mAbs can potently inhibit viral entry in monolayer cell lines overexpressing the viral receptor. This phenomenon is common among CoV mAbs, whereby the RBD is immunodominant and elicits potently neutralizing antibodies, making it a frequent target of CoV vaccine designs^28,29^.

### RBD mAb H501-008 and S2-specific mAb H501-101 cross-react with HCoV-HKU1 and - OC43

Because the donor’s serum reacted with both HCoV-HKU1 and HCoV-OC43 S proteins, we analyzed the cross-reactivity of the 14 isolated mAbs against S proteins from other beta-CoVs, including HCoV-OC43, MHV, MERS-CoV, SARS-CoV, and SARS-CoV-2. Interestingly, RBD-specific H501-008 and S2-specific H501-101 cross-reacted with cell surface-expressed HCoV-HKU1 and HCoV-OC43 S proteins, as well as soluble S proteins by ELISA **(Figures S4, S5A, and S5B)**. Neither antibody showed detectable cross-reactivity with the more distantly related MERS-CoV or sarbecovirus spikes tested **(Figure S4)**. The lack of cross-reactivity with epidemic CoVs was expected because the PBMCs were collected prior to the emergence of SARS-CoV-2, and the donor most likely had not been previously exposed to SARS-CoV or MERS-CoV.

We next performed surface plasmon resonance (SPR) experiments to more thoroughly characterize binding kinetics of H501-008 and H501-101 to HCoV-HKU1 and -OC43 RBD/CTD and S proteins. H501-008 Fab bound the HCoV-HKU1 RBD with subnanomolar affinity (161.0 pM) and also recognized the HCoV-OC43 CTD, albeit with reduced affinity (∼25 nM) that can primarily be attributed to an increased off-rate (**Figures S5C and S5D**). Similarly, H501-101 Fab bound HCoV-HKU1 and HCoV-OC43 with nearly equal affinities of 1.39 pM and 1.12 pM, respectively **(Figure S5E and S5F)**. The cross-reactive epitope recognized by the S2-specific mAb H501-101 is unsurprising given that the HCoV-HKU1 and HCoV-OC43 S2 share ∼72% amino acid similarity. Additionally, a plethora of cross-reactive CoV S2-specific mAbs have been isolated to date^30–32^. Beyond S2, these data allude to a cross-reactive epitope within the HCoV-HKU1 and -OC43 CTD, which is particularly intriguing because HCoV-HKU1 uses TMPRSS2 as its entry receptor, whereas HCoV-OC43 primarily engages 9-*O*-acetylated sialic acid through its S-NTD, for attachment and entry^14^.

### HCoV-HKU1/OC43 RBD mAb H501-008 protects against lethal OC43 challenge

We next tested the ability of H501-008 and H501-101 to neutralize authentic HCoV-OC43 and found that H501-008 weakly neutralized HCoV-OC43 (IC_50_ > 10 µg/mL with 60% neutralization shown at the highest mAb concentration tested, 40 µg/mL), whereas H501-101 did not neutralize the virus **(Figure S5G)**. To further assess whether the cross-reactive mAbs could confer protection in live virus challenge models, we intranasally infected mice with a neurovirulent strain of HCoV-OC43^9^ followed by intraperitoneal injection of the mAbs H501-008, H501-101, H501-022, or PBS **(Figure S5H)**. Infected mice were monitored daily for body weight loss, clinical disease score, and mortality. Mice injected with H501-101 or H501-022 succumbed to infection and showed progressive weight loss, severe clinical outcomes, and mortality similar to control animals **(Figure S5H, S5I, and S5J)**. In contrast, H501-008 completely protected the HCoV-OC43-challenged mice from severe disease, showing no signs of body weight loss, disease symptoms, and mortality **(Figure S5H, S5I, and S5J)**. Despite weak neutralization in monolayer cell culture, H501-008 conferred robust protection against lethal HCoV-OC43 challenge. There is 99.26 % sequence identity of HCoV-OC43 S RBD in the mouse model strain and live virus strain for which neutralization was tested; this discordance suggests that conventional *in vitro* neutralization assays may not fully capture antibody activity in physiologically relevant settings and raises the possibility that H501-008 interferes with viral entry or spread through mechanisms not reproduced in the monolayer assay.

### HCoV-HKU1/OC43 RBD mAb H501-008 binds outside the TMPRSS2-binding site and traps the RBD in the “up” conformation

Next, we sought to characterize the molecular determinants that mediate H501-008 cross-reactivity using cryogenic electron microscopy (cryo-EM) studies of H501-008 Fab in complex with HCoV-HKU1 S-2P **(Figure S6)**. We observed that H501-008 trapped the HCoV-HKU1 RBD in the rarely observed “up” conformation. Similar to RBD conformations observed in related betacoronaviruses, such as MERS-CoV and SARS-CoV^22,33,34^, the HCoV-HKU1 RBD projects upward from the rest of the S protein, exposing S2 heptad repeat 1 helices **(Figure 2A).** While we observed density corresponding to H501-008 Fabs bound to the “up” RBDs, conformational plasticity prevented us from solving the structure at high resolution. Thus, we performed X-ray crystallographic studies of H501-008 Fab bound to the isolated HCoV-HKU1 RBD and solved the complex structure to 2.60 Å resolution (**Table S2**). Despite this high resolution, the asymmetric unit contained only a single Fab–RBD complex, resulting in extremely high solvent content (∼70%) and relatively poor electron density in some regions of the map, which limited model building. Nevertheless, electron density at the interface between the H501-008 CDRs and the HCoV-HKU1 RBD was interpretable, and the Fab binding orientation agreed well with that observed in the lower-resolution cryo-EM map **(Figures 2A and 2B).**

**Figure 2.**
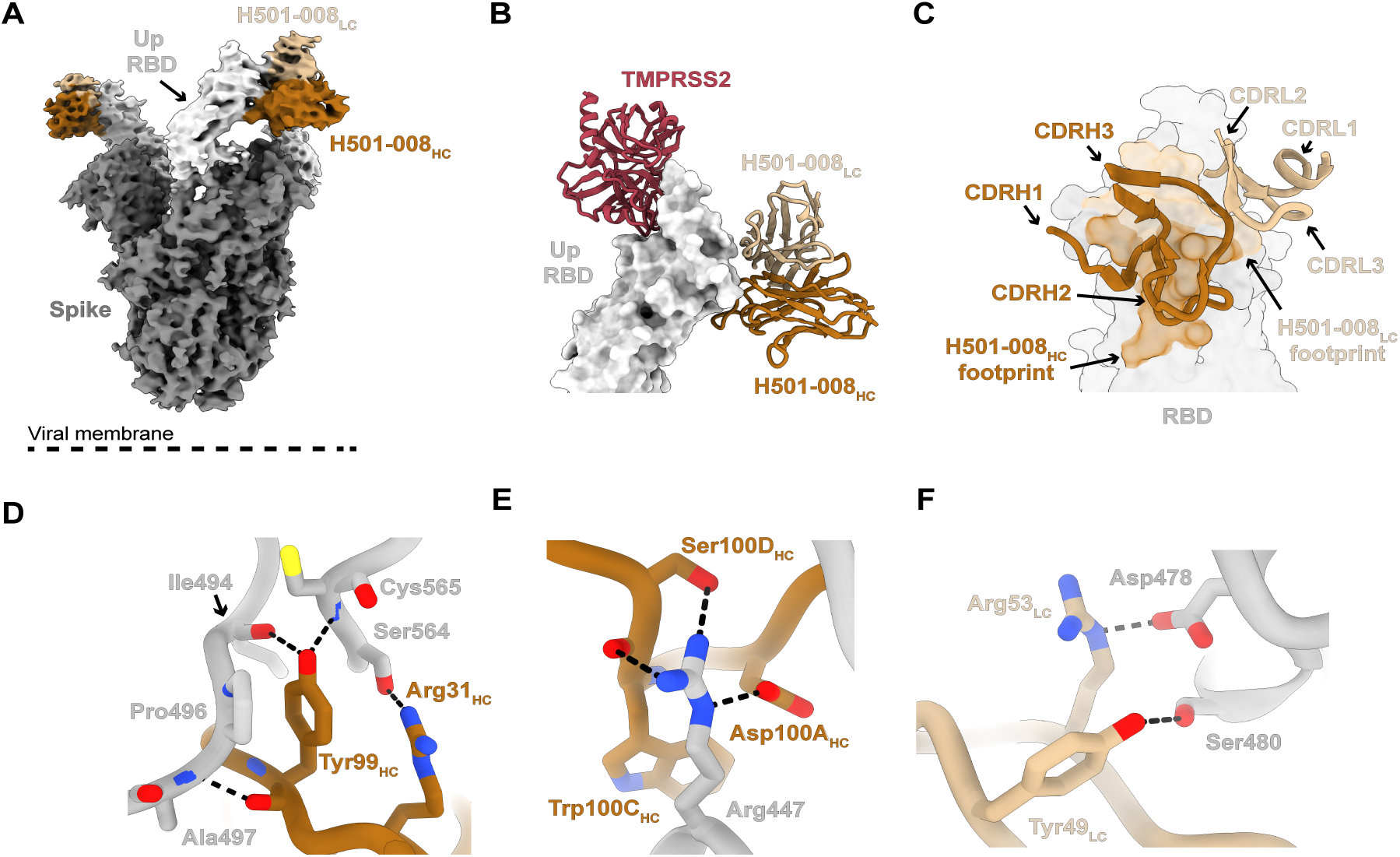
Structure of HCoV-HKU1 S-2P bound to H501-008. **(A)** Cryo-EM map of HCoV-HKU1 S-2P bound to H501-008. **(B)** Superimposition of the TMPRSS2-RBD structure (PDB 8Y8B) by alignment on the RBD bound to the H501-008 V_H_ and V_L_ domains from the crystal structure. The RBD is shown as a surface representation, whereas TMPRSS2 and H501-008 are displayed as ribbons. **(C)** Close-up view of H501-008-mediated contacts on the RBD epitope, colored by V_H_ and V_L_ domain contributions. **(D,E)** Interacting residues between the H501-008 heavy chain and the HCoV-HKU1 C RBD. **(F)** Interacting residues between the H501-008 light chain and the HCoV-HKU1 RBD. Residues are shown as sticks. Colors are as follows: S (gray), RBD (light gray), H501-008 V_H_ (orange), H501-008 V_L_ (tan), TMPRSS2 (red). Potential hydrogen bonding indicated by black dashed line.

Analysis of the hybrid structural data revealed several key contacts distributed across the CDRH1, CDRH3, and CDRL2 loops **(Figure 2C).** Key contacts include potential hydrogen bonds between Arg31_HC_ and the side chain of Ser564, as well as between the backbone amine of Cys565 and Tyr99_HC_, which simultaneously interacts with the backbone carboxyl of Ile494 **(Figure 2D)**. In addition to Tyr99_HC_, CDRH3 residues Asp100_A-HC_ and Ser100_D-HC_ form hydrogen bonds with Arg447 via their side chains **(Figure 2E)**. H501-008 light-chain interactions are largely confined to CDRL2, where Tyr49_LC_ and Arg53_LC_ form hydrogen bonds with the side chains of Ser480 and Asp478, respectively **(Figure 2F)**.

By comparing key contact residues within the HCoV-HKU1 RBD to their counterparts in the HCoV-OC43 RBD, we began to define the molecular determinants that permit H501-008 cross-reactivity **(Figure S7)**. HCoV-HKU1 RBD shares 53.2% sequence identity with HCoV-OC43 RBD. Despite this modest overall sequence conservation, several key residues that mediate H501-008 binding, including Arg447 and Ala497, are strictly conserved in HCoV-OC43 as Arg463 and Ala524, respectively **(Figure S7)**. The diminished binding affinity observed for HCoV-OC43 RBD may result from an insertion present in HCoV-HKU1 RBD at positions 558–565. Although additional contact residues differ between the two RBDs (e.g., Ile494), these variations likely have a smaller impact on binding because the corresponding H501-008 residues interact primarily with the peptide backbone, which remains structurally invariant despite side-chain differences.

To further define the H501-008 epitope, we performed alanine-scanning mutagenesis of HCoV-HKU1 and HCoV-OC43 S proteins followed by cell-surface mAb-binding assays. These studies revealed that Arg447 and Ala497 are critical for H501-008 binding to HCoV-HKU1 S **(Figure S8A)**. Similarly, Arg463 and Ala524 were important for H501-008 binding to HCoV-OC43 S **(Figure S8B)**. HCoV-HKU1 pseudovirus neutralization assays using S mutants further confirmed the cell-surface binding data, emphasizing the importance of Ala497 in the H501-008 interaction with HCoV-HKU1 S **(Figure S8D).** The R447A mutant pseudovirus exhibited reduced infectivity in HEK293T-TMPRSS2 cells, potentially due to conformational changes in the S RBD **(Figure S8C)**. Together, these structural and mutational data define a conserved cross-reactive epitope outside the TMPRSS2-binding site.

### HCoV-HKU1 RBD mAb H501-018 recognizes RBD in two conformational states

To define the model of antibody recognition within the major RBD competition group (**Figure 1**), we determined a 3.4 Å resolution single-particle cryo-EM structure of HCoV-HKU1 S-2P bound to H501-018 Fab. We identified two distinct conformational states of the H501-018 Fab– S-2P complex. In State 1, two RBDs adopt the “up” conformation, whereas one remains “down” **(Figure S6B)**. In State 2, only a single RBD adopts the “up” conformation **(Figure 3A)**. To better resolve molecular interactions at the Fab–RBD interface, we performed local classification and refinement of cryo-EM density corresponding to H501-018 Fab bound to a “down” RBD. This approach yielded a locally refined density map at 3.2 Å resolution, allowing us to model the H501-018–RBD complex **(Figure 3A)**^35^. The structure revealed a binding orientation for H501-018 that engages an RBD epitope outside the TMPRSS2 receptor-binding motif **(Figure 3B)**, suggesting that neutralization does not occur through direct competition with TMPRSS2.

**Figure 3.**
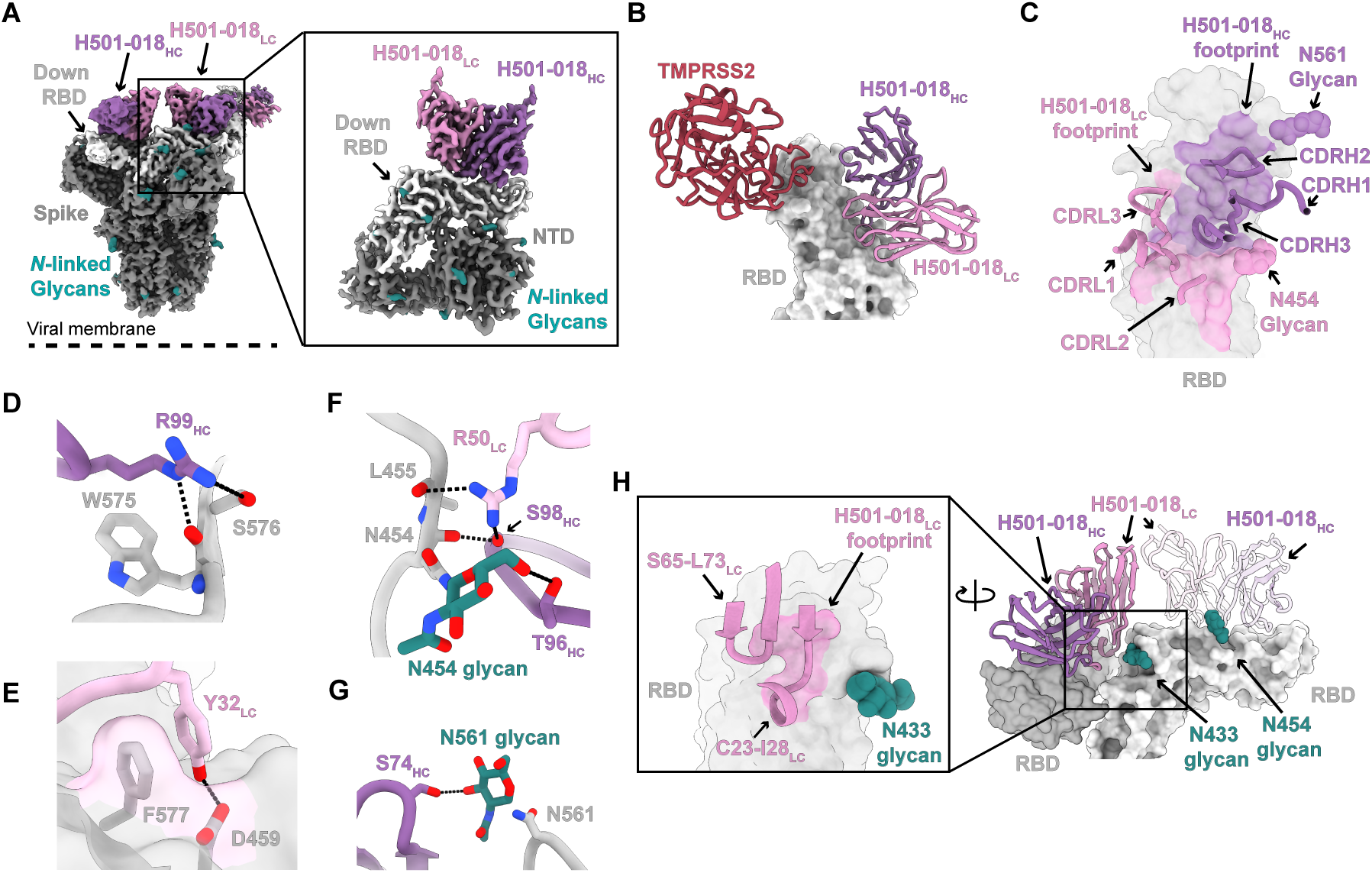
Structure of HCoV-HKU1 S-2P bound to H501-018. **(A)** Cryo-EM map of HCoV-HKU1 S-2P bound to H501-018 (State 2; see also Figure S6). Inset shows the locally refined map of the V_H_ and V_L_ domains, the RBD, and the NTD. **(B)** TMPRSS2 (PDB 8Y8B) is aligned based on the RBD bound to H501-018 V_H_ and V_L_ from the locally refined cryo-EM structure. RBD is displayed as a surface representation, and TMPRSS2 and H501-018 are displayed as ribbons. **(C)** Close up view of the H501-018 mediated contacts on the RBD epitope colored by the V_H_ and V_L_ domain contributions. **(D-G)** Interacting residues between H501-018 and HCoV-HKU1 RBD. Residues are shown as sticks. **(H)** H501-018 bound to a “down” RBD in State 2 also contacts the adjacent RBD protomer, forming a quaternary component of the epitope. Inset shows the RBD epitope from the neighboring H501-018 light chain. Colors are as follows: S (gray), RBD (light gray), H501-018 V_H_ (purple), H501-018 V_L_ (pink), *N*-linked glycans (teal), TMPRSS2 (red). Potential hydrogen bonding indicated by black dashed line.

Analysis of the Fab–RBD interface revealed contacts involving all H501-018 CDR loops and an epitope buried surface area (BSA) of 1,064 Å^2^ (551 Å^2^ against the heavy chain and 513 Å2 against the light chain) **(Figures 3B and 3C)**^36^. Critical contacts involve a combination of germline-encoded and somatically hypermutated H501-018 residues, including Arg99_HC_, which forms hydrogen bonds with the backbone carbonyl of Trp575 and the side chain of Ser576 (**Figure 3D**). Tyr32_LC_ within CDRL1 forms a π-stacking interaction with Phe577 and a hydrogen bond with Asp459 in the RBD **(Figure 3E)**. In addition, Arg50_LC_ forms a hydrogen-bonding network at the base of the Asn454 glycan **(Figure 3F)**. Ser98_HC_ also forms a hydrogen bond with the backbone carbonyl of Asn454, an interaction that may have been hindered, had Ser98 not arisen through somatic hypermutation from a previous Trp98_HC_^37,38^. We also observed two glycan-mediated interactions between the heavy chain and the RBD. Thr96_HC_ of the CDRH3 loop forms a hydrogen bond with the Asn454 glycan **(Figure 3F)**, an interaction enabled by the W98S_HC_ somatic mutation that accommodates glycan recognition. A second glycan-mediated interaction occurs between the Asn561 glycan and Ser74_HC_ **(Figure 3G)**. Interestingly, in the State 2 complex, framework regions 1 and 3 of H501-018 engage the “down” RBD of the neighboring protomer, establishing a quaternary epitope that likely contributes to increased Fab affinity at the trimer interface in this state **(Figure 3H)**.

### H501-022 targets the TMPRSS2-binding site and blocks receptor engagement

To elucidate the molecular mechanism by which H501-022 recognizes the RBD and blocks HCoV-HKU1 pseudovirus entry, we determined a 3.6 Å resolution single-particle cryo-EM structure of HCoV-HKU1 S-2P bound to H501-022 Fabs **(Figure 4A, Figure S6C, Table S3)**. Global maps revealed all three RBDs adopting the “up” conformation and bound to H501-022 Fab **(Figure 4A)**. Given the structural plasticity of the “up” conformation, we performed local classification and refinement at the Fab–RBD interface to generate cryo-EM maps suitable for model building and refinement **(Figure 4B, Figure S6C, Table S3)**. H501-022 directly overlaps the RBD–TMPRSS2 receptor-binding site, providing a structural mechanism for its potent neutralization through inhibition of viral engagement with cell-surface TMPRSS2 **(Figure 4B)**. While all CDR loops engage the epitope, the majority of interactions are contributed by the CDRH2 and CDRH3 loops, which account for 31.8% and 21.5% of the total paratope BSA, respectively **(Figure 4C)**^36^. Although both chains contribute to the interaction, the light chain engages the TMPRSS2-binding site to a lesser extent than the heavy chain **(Figure 4C)**. Within the light-chain interface, CDRL3 residues engage a patch on the RBD centered around Asn531 and Lys536. Tyr92_LC_ forms hydrogen bonds with both the backbone and side chain of Asn531, while nearby residues Ser93_LC_ and Ser94_LC_ contribute additional hydrogen bonding through backbone contacts (**Figure 4D**). Tyr32_LC_ and Arg30_LC_ further stabilize the interaction by contacting Lys536 and Lys537, respectively **(Figure 4E)**.

**Figure 4.**
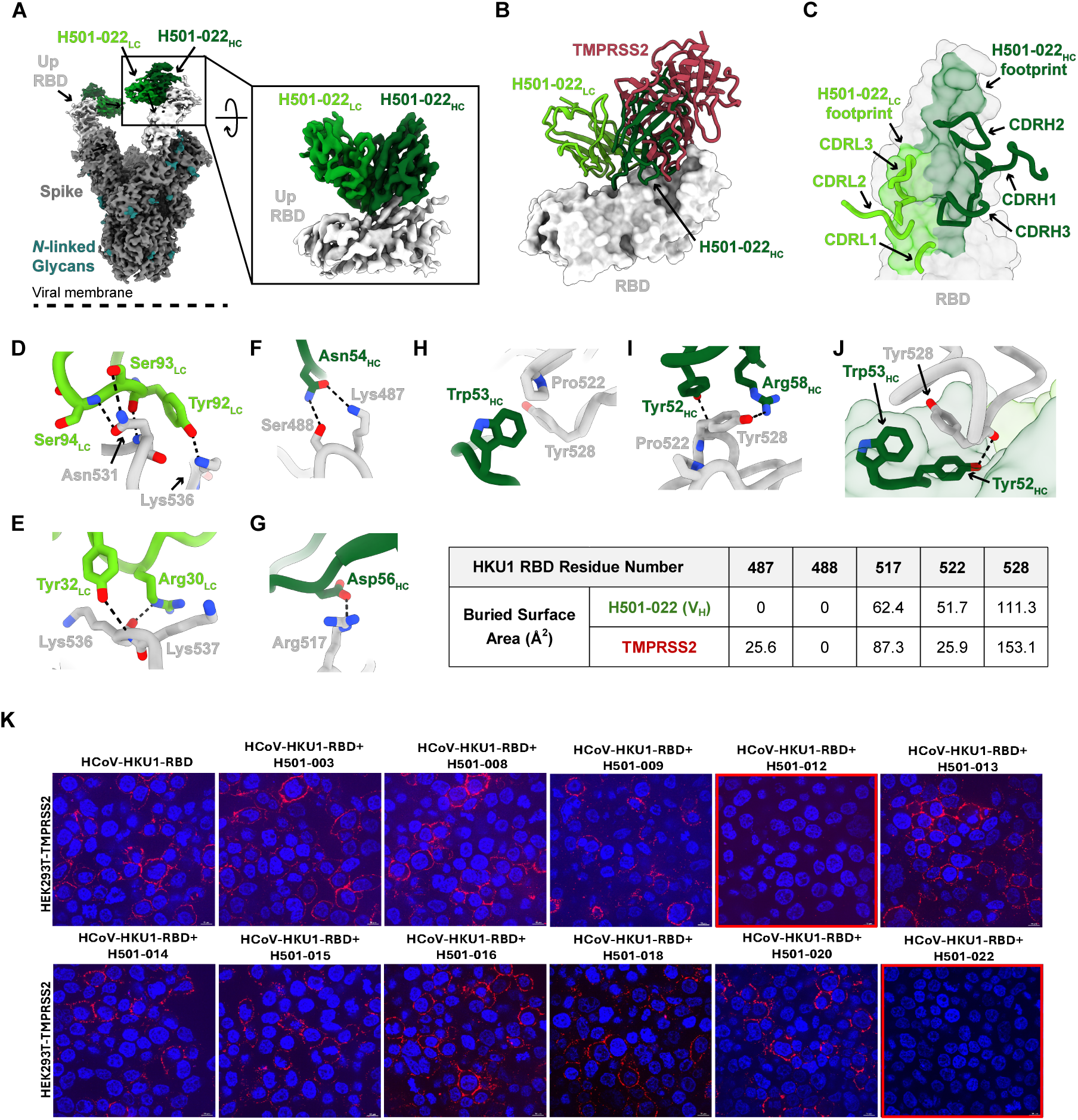
Structure of HCoV-HKU1 S-2P bound to H501-022. **(A)** Cryo-EM map of HCoV-HKU1 S-2P bound to H501-022. Inset shows the locally refined map of the V_H_ and V_L_ domains, and the S1 RBD.**(B)** Superimposition of the TMPRSS2-RBD structure (PDB 8Y8B) by alignment on the RBD bound to H501-022 V_H_ and V_L_ from the locally refined cryoEM structure. **(C)** Close up view of the H501-022 mediated contacts on the RBD epitope colored by the V_H_ and V_L_ domain contributions. H501-022 CDR loops are shown as ribbons, and the RBD is a surface representation. **(D-E)** Interacting residues between H501-022 light chain and HCoV-HKU1 RBD. **(F-J)** Interacting residues between H501-022 heavy chain and HCoV-HKU1 RBD. The table shows the Buried Surface Area (BSA) for residues on the HCoV-HKU1 RBD contributed by H501-022 heavy chain or TMPRSS2. Colors are as follows: S (gray), RBD (light gray), H501-022 V_H_ (green), H501-022 V_L_ (light green), *N*-linked glycans (teal), TMPRSS2 (red). Potential hydrogen bonding indicated by black dashed line. (**K**) HCoV-HKU1 RBD mAbs were analyzed for inhibition of S-TMPRSS2 interaction by immunostaining of cell surface-bound HCoV-HKU1 RBD-mAb complexes using confocal microscopy. The red highlighted box denotes mAbs that block S-TMPRSS2 interaction.

The CDRH2 loop plays a critical role in blocking TMPRSS2. It engages several RBD residues previously identified as TMPRSS2 contact sites, including Lys487, Arg517, Pro522, and Tyr528 **(Figures 4F–4J)**^39,40^. Asn54_HC_ forms hydrogen bonds with Lys487 and Ser488, while Asp56_HC_ forms a hydrogen bond with Arg517 **(Figures 4F and 4G)**. Trp53_HC_ stacks against Pro522, while Tyr52_HC_ and Arg58_HC_ form hydrogen bonds with Tyr528 **(Figures 4H–4J)**. Importantly, Tyr528 lies within a pocket at the V_H_–domain interface, contributing 111 Å^2^ of BSA **(Figure 4J)**. Supporting the structural data, cell-surface binding assays showed that H501-022 blocks HCoV-HKU1 S binding to cell-surface-expressed TMPRSS2 **(Figures 4K and S9A),** and this extensive interaction with Tyr528 helps explain why mutating Tyr528 to alanine substantially reduced H501-022 binding to cell-surface-expressed HCoV-HKU1 S **(Figure S9B)**.

We next compared H501-022 with HCoV-HKU1 RBD-specific antibody R7, a recently reported HCoV-HKU1 RBD-targeting mAb^21^ that was isolated by phage display and also binds the S–TMPRSS2 interface. Because R7 pseudovirus neutralization had not been previously reported, we compared the cell-surface binding and pseudovirus neutralization properties of R7 and H501-022 against HCoV-HKU1 genotypes A and C. Both R7 and H501-022 bound HCoV-HKU1 genotype A S protein, whereas R7 failed to bind genotype C **(Figure S10A).** Consistent with these results, both antibodies neutralized the entry of genotype A pseudovirus; however, only H501-022 neutralized genotype C **(Figure S10B)**. Although H501-022 and R7 target overlapping epitopes that coincide with the TMPRSS2-binding site **(Figure S10C–N),** differences in their interactions with the RBD may contribute to their distinct genotype breadth. In particular, residues 530, 532, 536, and 538 contribute substantially to the H501-022 interface with the genotype C RBD but make little to no contribution to the R7 interface with the genotype A RBD **(Figure S10N)**. These differences in epitope engagement may help explain why H501-022 neutralizes both genotypes A and C, whereas R7 neutralizes only genotype A. Furthermore, multiple sequence alignment of HCoV-HKU1 S protein sequences from genotypes A, B, and C demonstrates that critical H501-022 contact residues are well conserved across all HCoV-HKU1 strains **(Figure S11),** consistent with H501-022 being a broadly reactive HCoV-HKU1 mAb targeting the S– TMPRSS2 interface. Collectively, these structures define three distinct modes of HKU1 RBD recognition that differ in receptor competition, conformational accessibility, genotype breadth, and cross-reactivity, and therefore, provide a molecular framework for understanding neutralizing antibody recognition of endemic beta-HCoVs **(Figure S12**).

### HCoV-HKU1 RBD mAbs do not potently neutralize live virus infection in primary human airway epithelial cells

Prior to the description of TMPRSS2 as a functional receptor, HCoV-HKU1 live virus infection could only be measured in primary airway epithelial cells differentiated at an air–liquid interface^41^. More recently, it has become standard to perform HCoV-HKU1 pseudovirus neutralization assays in TMPRSS2-overexpressing cell lines, as we described above. However, another report of HCoV-HKU1-specific mAbs showed varying neutralization capacity in primary human airway epithelial cells^42^. Thus, we set out to assess the neutralization capacity of H501-008, -018, and -022 against live HCoV-HKU1 in differentiated normal human bronchial epithelial (NHBE) cells, a physiologically relevant cell culture system.

First, to determine whether the three RBD mAbs exhibit comparable binding to the S from HCoV-HKU1-B used in the live-virus assays relative to the HCoV-HKU1-C S, we evaluated binding to cell surface-expressed S. H501-018 and H501-022 showed similar levels of HCoV-HKU1-B S binding compared to HCoV-HKU1-C S, whereas H501-008 showed slightly lower binding to HCoV-HKU1-B S **(Figure S13A)**. Amino acid A497 mutated to T487 in the HCoV-HKU1-B strain used in our assays; this residue is critical for H501-008 binding **(Figure S8A and S8D)**. In addition, we performed an HCoV-HKU1-B pseudovirus neutralization assay with the three RBD mAbs; H501-022 and -018 showed potent pseudovirus neutralization on par with HCoV-HKU1-C, and, as expected, H501-008 showed reduced neutralization against HCoV-HKU1-B **(Table 1, Figure S13B)**.

We collected apical washes from NHBE cells following infection with HCoV-HKU1 preincubated with mAbs and quantified HCoV-HKU1 RNA by RT-qPCR at 24 and 48 hours post-infection. Surprisingly, none of the HCoV-HKU1 RBD mAbs potently inhibited live HCoV-HKU1 infection **(Figure 5A–5C)**, in contrast to their activity in pseudovirus assays **(Figure S13B)**. H501-018 and H501-022 exhibited neutralization at only the highest concentration tested (10 µg/mL), whereas H501-008 showed no detectable neutralization **(Figure 5A–5C)**. Western blot analysis of HCoV-HKU1 N protein in infected NHBE cells yielded similar results, showing minimal inhibition of viral infection by the RBD mAbs **(Figure 5D)**. We observed the same pattern in HCoV-HKU1 neutralization assays using NHBE cells from a different donor **(Figure S14)**.

**Figure 5.**
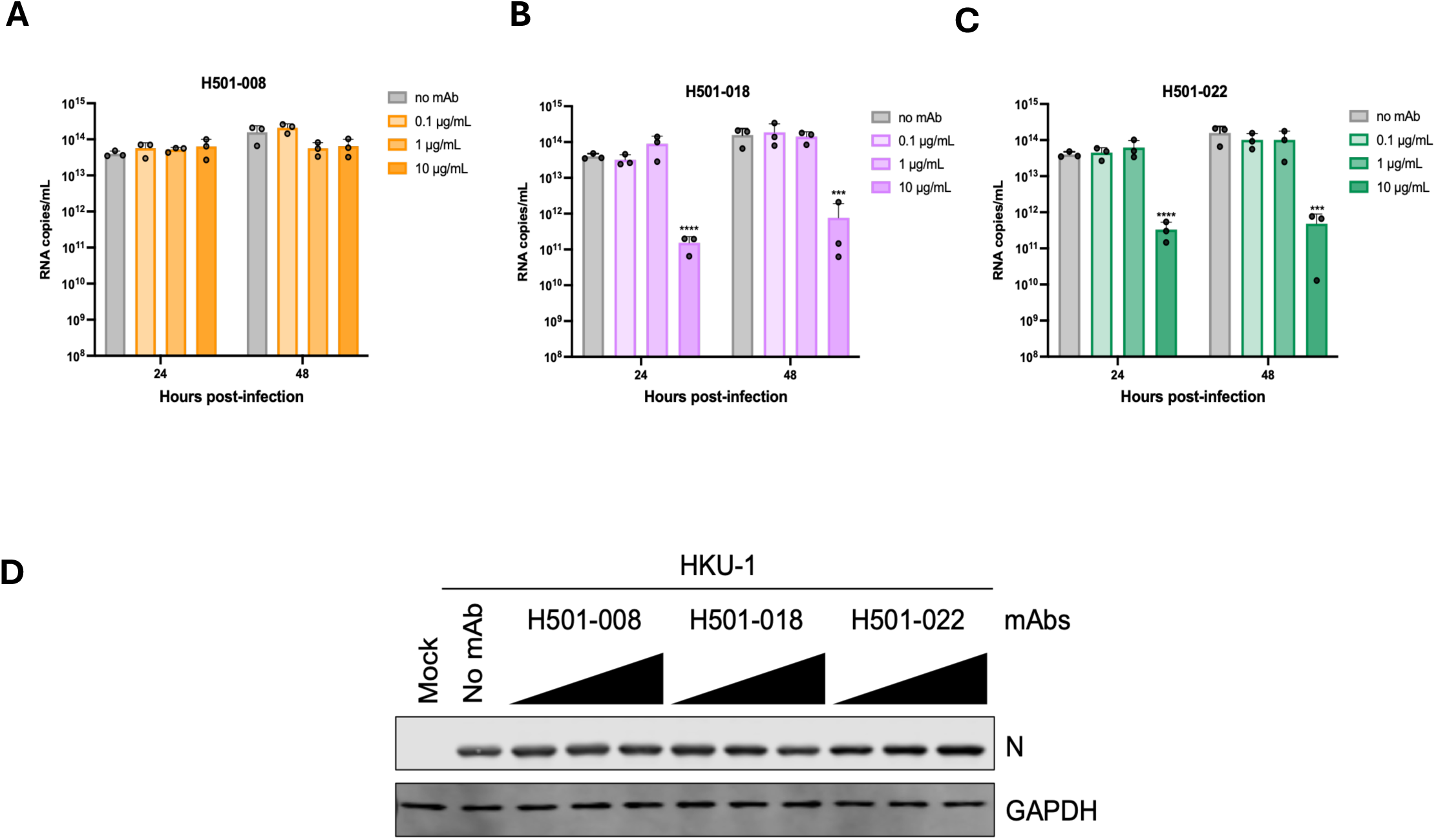
HCoV-HKU1 live virus neutralization with CTD mAbs. Concentration-dependent neutralization effect of CTD mAbs **(A)** H501-008, **(B)** H501-018, and **(C)** H501-022 against HCoV-HKU1 live virus infection in NHBE cells (Donor 1) was analyzed by RT-PCR of viral nucleocapsid (N) gene. Twenty-four and 48 hours post-infection, RNA copies/ ml was calculated with reference to the standard curve generated using in vitro–transcribed RNA corresponding to the HKU1 N gene. Error bars denote mean ± SD. RNA copies at Individual time points were compared to no mAb control using one-way ANOVA with Dunnett’s multiple comparison. ***P ≤ 0.001, ****P ≤ 0.0001. **(D)** HCoV-HKU1 was treated with increasing concentrations of mAbs prior to infection, and cell lysates collected at 96 hours post-infection were analyzed by Western blotting to assess viral N protein expression. GAPDH was used as an internal control.

These findings demonstrate that potent pseudovirus neutralization in TMPRSS2-overexpressing monolayer cells does not necessarily predict neutralization of authentic HCoV-HKU1 infection in primary human airway epithelial cells, demonstrating the importance of assessing CoV mAb neutralization efficacy in physiologically relevant models. Moreover, the weak neutralization exhibited by H501-022, despite blocking S interactions with TMPRSS2, suggests the possibility of TMPRSS2-independent viral entry in NHBE cells. Further research should explore the neutralization potential of the RBD mAbs in diverse airway epithelial cell models, such as upper and lower airway cells, which might reveal viral tropism and varied receptor usage. Since these results challenge current understanding regarding RBD-specific antibody neutralization potency, further characterization of HCoV-HKU1 neutralization-sensitive antibody epitopes is needed.

## Discussion

HCoVs represent a diverse group of respiratory viruses with varied clinical impacts^43^. While epidemic CoVs, such as SARS-CoV-2, have caused deadly outbreaks, four endemic HCoVs, HCoV-229E, -NL63, -OC43, and -HKU1, circulate globally and typically cause mild respiratory disease, particularly in children. Elderly, immunocompromised, and otherwise co-morbid individuals are at higher risk for severe HCoV disease^44^. Despite their broad circulation and contribution to the global disease burden, endemic HCoVs remain understudied, particularly with respect to vaccine and therapeutic development. This gap is concerning for several reasons. First, HCoV co-infection with other respiratory viruses can affect disease severity^3^. Additionally, HCoV infections may influence immune imprinting throughout life. Moreover, HCoV prevalence may shape the emergence potential of other CoVs.

Herein, we address a critical gap in our understanding of humoral immunity to endemic HCoVs by isolating and characterizing mAbs from a pre-COVID-19-pandemic HCoV-HKU1 convalescent donor. The 14 identified mAbs recognize diverse antigenic sites on HCoV-HKU1 S, the key surface glycoprotein responsible for receptor binding and membrane fusion. Structural analyses revealed three distinct antibody footprints within and around the TMPRSS2 receptor-binding site, a region essential for entry into host cells **(Figure S12)**. Importantly, RBD-specific mAbs H501-012 and H501-022 exhibited potent pseudovirus neutralization by directly blocking TMPRSS2 engagement, a mechanism reminiscent of SARS-CoV-2 Class 1 RBD mAbs that bind the “up” RBD conformation and block S–ACE2 receptor interactions^34,45^. Recently, another group described an HCoV-HKU1 RBD-specific mAb (R7) identified through a phage-display platform that overlaps with the TMPRSS2-binding motif. However, that study used the recombinant RBD instead of the full-length S for structural characterization and therefore did not provide information on recognition of “up” and “down” RBD conformations. Additionally, the ability of R7 to neutralize HCoV-HKU1 was not determined^21^. We compared HCoV-HKU1 mAb H501-022 with R7 for pseudovirus neutralization and S-binding assays and found that R7 could only bind and neutralize HCoV-HKU1 genotype A, whereas H501-022 bound and neutralized both genotypes A and C. The data show the broader reactivity of H501-022 to multiple genotypes of HCoV-HKU1.

Characterization of H501-008 revealed potent pseudovirus neutralization of HCoV-HKU1 and significant cross-reactivity against HCoV-OC43, mapping antibody binding to a conserved RBD epitope that may confer broader HCoV recognition. H501-008-like mAbs resemble Class V SARS-CoV-2 RBD-specific antibodies that bind exclusively to the “up” RBD conformation and exhibit broad sarbecovirus reactivity^46^. Remarkably, despite H501-008 showing relatively weaker neutralization of HCoV-OC43 in monolayer cells, it protected against lethal HCoV-OC43 infection in a mouse model, suggesting the potential role of HCoV-OC43 CTD in viral entry in physiologically relevant systems. In contrast, H501-018, representative of the predominant RBD-specific competition group, neutralized HCoV-HKU1 less potently but engaged both “up” and “down” RBD conformations, reminiscent of SARS-CoV-2 Class 2 anti-RBD antibodies^34,45^. The distinct epitope targeted by H501-018, which is separate from both the TMPRSS2- and H501-008-binding sites, highlights the presence of multiple antigenic surfaces within the HCoV-HKU1 RBD, as with HCoV-OC43, capable of eliciting neutralizing responses.

Indeed, identification of mAbs (H501-012 and H501-022) that potently inhibit HCoV-HKU1 pseudovirus entry by engaging the TMPRSS2 receptor-binding motif further highlights the immunological vulnerability of this region. Analogous to SARS-CoV-2 antibodies that block ACE2 engagement and potently neutralize infection^45,47^, our findings suggest a parallel neutralization mechanism in an endemic beta-CoV. Notably, antibodies recognizing non-RBD regions also contribute to SARS-CoV-2 neutralization and are particularly important because the elevated mutation rate within the RBD renders RBD-specific antibody responses susceptible to viral escape^48–50^. Similarly, identification of mAbs targeting diverse regions of HCoV-HKU1 S beyond the RBD extends this concept to endemic CoVs and supports TMPRSS2-binding site-adjacent regions as potential targets for broader CoV vaccine design. Although the NTD- and S2-directed mAbs isolated in this study did not neutralize HCoV-HKU1 pseudovirus, both NTD- and S2-specific mAbs have been shown to mediate neutralization in other CoVs^51,52^. An unexpected finding of this study was the marked difference between pseudovirus and authentic virus neutralization. In contrast to the pseudovirus neutralization observed with the RBD mAbs, H501-018 and H501-022 inhibited live virus entry only at the highest concentration tested whereas H501-008 did not inhibit live virus entry. These findings suggest that, although RBD mAbs potently neutralize HCoV-HKU1 pseudovirus in a TMPRSS2-overexpressing cell line, this activity does not fully translate to authentic virus infection in physiologically relevant primary airway epithelial cells. These observations also raise the possibility that antibodies targeting non-RBD epitopes may reveal novel mechanisms of viral neutralization. Moreover, the sub-neutralizing nature of H501-022, despite blocking S interaction with TMPRSS2, suggests that additional host factors could contribute to HCoV-HKU1 infection in airway cells. Future studies evaluating neutralization in both upper and lower airway cells may further define tropism-specific contributions of TMPRSS2 and other host factors to HCoV-HKU1 entry.

Identification of H501-008 as an HCoV-OC43-cross-reactive mAb reemphasizes the existence of conserved antigenic epitopes among beta-CoVs. Notably, H501-008 protected against lethal HCoV-OC43 challenge in a mouse model, despite showing only weak neutralization in monolayer cell culture. Together with the discordance observed between HKU1 pseudovirus and authentic-virus neutralization, this finding further demonstrates that conventional in vitro neutralization assays may not fully predict antibody activity in more physiologically relevant settings. Furthermore, discovery of cross-reactive mAbs has become a major focus of recent CoV research^53,54^. Several studies have described broadly neutralizing mAbs isolated from SARS-CoV-2 survivors that cross-react with SARS-CoV and other beta-CoVs^55–58^. In addition, mAbs targeting the S2 stem helix, which is highly conserved across all known HCoVs^59–62^, and fusion peptide-targeting mAbs exhibit cross-neutralization across all known HCoVs^63^. These findings suggest that defining antibody recognition across the intact spike can reveal conserved and conformational epitopes that may be missed when focusing exclusively on isolated subdomains, supporting the use of full-length S trimer vaccine antigens.

While recent mAb development efforts have largely focused on epidemic CoVs, this study provides an important perspective on less-appreciated endemic strains. Although endemic HCoVs are often associated with less severe disease outcomes, they continue to cause upper respiratory tract infections, hospitalization, and life-threatening illness in high-risk populations^3–6^. Further, understanding the fundamental biology and immunology of endemic HCoVs that, after evolving with humans over time, may also provide insight into how newly emerged coronaviruses adapt to sustained circulation in humans, as is now occurring with SARS-CoV-2. The absence of licensed vaccines or mAb therapies targeting endemic HCoVs reflects both a gap in research attention and an opportunity to better define the principles governing antibody-mediated protection against these viruses.

In conclusion, isolation of mAbs targeting diverse epitopes on the HCoV-HKU1 S protein, including epitopes within and around the TMPRSS2 receptor-binding site, expands our understanding of the antigenic landscape of endemic HCoVs. The structural characterization presented here, together with the differences in HCoV-HKU1 neutralization observed in receptor-expressing monolayer cells and primary airway epithelial cells, reveals that structural epitope definition and neutralization measured in simplified cell systems must be interpreted in the context of physiologically relevant models. Further research should investigate the neutralization potential of the RBD-specific mAbs in diverse airway epithelial cell models, including upper and lower airway cells, which may reveal differences in viral tropism and receptor usage. Because these results challenge the assumption that potent receptor-blocking activity in conventional neutralization assays predict activity in primary airway epithelium, future studies should define additional neutralization-sensitive epitopes and mechanisms using physiologically relevant models. Together, this work defines distinct modes of antibody recognition within and around the HKU1 receptor-binding site and demonstrates that the biological context of viral entry is a critical determinant of antibody activity against endemic coronaviruses that will be essential for improving pandemic preparedness and developing durable protection against evolving CoVs.

## Methods

### Cell Lines

HEK293T (ATCC CRL-3216) cells were maintained in Dulbecco’s modified Eagle’s medium (DMEM) supplemented with 10% FBS, 2 mM glutamine, and 1% penicillin-streptomycin. Huh7 cells were kindly provided by Dr. Mark Heise at the University of North Carolina at Chapel Hill and were grown in DMEM supplemented with 10% FBS and anti/anti. Expi293F cells (#A14527, Thermo Fisher Scientific) were maintained in Expi293 expression medium at 37°C, 8% CO_2_, and 70% humidity with shaking at 200 rpm. FreeStyle293-F cells (#R79007, Thermo Fisher Scientific) were maintained in FreeStyle293 expression medium at 37°C, 8% CO_2_, and 70% humidity with shaking at 125 rpm.

### Human PBMCs and viral isolates

Serum and PBMC samples from an individual following a confirmed HCoV-HKU1 infection were collected under the National Institutes of Health, National Institute of Allergy and Infectious Diseases, Vaccine Research Center (NIH, NIAID, VRC) clinical protocol VRC 200 (ClinicalTrials.gov: NCT00067054). All study procedures were reviewed and approved by the Institutional Review Boards (IRBs) of the NIH and NIAI. Participants received compensation for their time and inconvenience. Before ELISA testing, aliquots of frozen serum samples were heat-inactivated at 56°C for 1 h, and assays were conducted in at least two technical replicates. Cryopreserved PBMCs were thawed in a 37°C water bath and immediately transferred to ice-cold staining buffer for downstream processing. HCoV-HKU1-B used in live virus neutralization was isolated in a protocol approved by the IRB. Written informed consent was obtained from all participants before enrollment.

### DNA and protein vector constructs

Full-length S genes from HCoV-HKU1-C (GenBank ID: DQ339101), HCoV-HKU1-B (sequenced herein), HCoV-OC43 (GenBank ID: AIL49484.1), SARS-CoV (GenBank ID: AAP13441.1), SARS-CoV-2 (GenBank ID: QHD43416.1), MERS-CoV (GenBank ID: AFS88936.1), and MHV A59 (GenBank ID: AF029248.1) were cloned into mammalian expression vector VRC8400^64,65^ and confirmed by sequencing. S-2P, S1, and RBD were synthesized by PCR using full-length S templates, cloned into plasmid vector VRC8400, and sequence confirmed for protein production. For the expression of HCoV-HKU1 RBD-hFc, RBD residues (326–605) were fused C-terminally to the Fc domain of human immunoglobulin (Ig) previously cloned into a eukaryotic expression vector pCDNA3.1 (GenScript). Proteins used for ELISA binding contained a C-terminal 6× His-tag for protein purification. Genes encoding S ectodomains of HCoV-HKU1 (residues 1–1,276), HCoV-OC43 (residues 1–1,286), SARS-CoV (residues 1–1,190), SARS-CoV-2 (residues 1–1,208), and MERS-CoV (residues 1–1,291) used to generate S-2P proteins have been described previously^11,12,22^. S-2P constructs used for ELISA binding contained a C-terminal T4 fibritin trimerization motif, an HRV3C protease cleavage site, a Twin StrepTag, and an 8x His-tag for protein purification. Constructs used for generating S-based probes contained an Avi-tag for biotinylation. Proteins were expressed by transfecting vectors into Expi293F cells as described previously^28^. Transfected cell culture supernatants were collected and purified using HisTrap HP Hiload 16/60 Superdex columns (GE Healthcare).

### Mammalian protein expression and purification

Mammalian expression plasmids encoding HCoV-HKU1 RBD (residues 310–606 of isolate N5), HCoV-OC43 S1-CTD (residues 318–608), HCoV-HKU1 NTD (residues 1–294 of isolate N5), HCoV-HKU1 S (residues 1–1,276 of isolate N5), HCoV-OC43 S (residues 1–1,286), the heavy chain of H501-008 with an HRV3C cleavage site in the hinge between the CH1 and CH2 domains, the light chain of H501-008, the heavy chain of H501-101 with an HRV3C cleavage site in the hinge between the CH1 and CH2 domains, and the light chain of H501-101 were transfected into FreeStyle 293-F cells using polyethylenimine (PEI). Proteins containing a C-terminal HRV3C cleavage site, an Fc-tag, and an 8× His-tag (HCoV-HKU1 RBD, HCoV-OC43 S1-CTD, and HCoV-HKU1 NTD) and IgGs (H501-008 and H501-101) were purified using Protein A resin (Pierce Biotechnology). HCoV-HKU1 S-2P protein, containing a C-terminal HRV3C cleavage site, a T4 fibritin trimerization motif, an 8× His-tag and a Twin-Strep-tag, was purified using StrepTactin resin (IBA Lifesciences). Affinity-purified HCoV-HKU1 RBD, HCoV-OC43 S1-CTD, HCoV-HKU1 NTD, H501-008 IgG, and H501-101 IgG were purified by size-exclusion chromatography using a Superdex 200 Increase column (GE Healthcare) in 2 mM Tris pH 8.0, 200 mM NaCl, and 0.02% NaN3. Affinity-purified HCoV-HKU1 and -OC43 S-2P were purified by size-exclusion chromatography using a Superose 6 column (GE Healthcare) in 2 mM Tris pH 8.0, 200 mM NaCl, and 0.02% NaN3. H501-008 and H501-101 IgGs were digested by cleavage with 5% (wt/wt) HRV3C protease on ice for 4 h. Cleaved protein was passed over Protein A resin and Ni-NTA resin (Thermo Fisher Scientific) to remove cleaved Fc and excess protease. H501-008 and -101 Fabs were then purified by size-exclusion chromatography using a Superdex 200 Increase column in 2 mM Tris pH 8.0, 200 mM NaCl, and 0.02% NaN3.

### Isolation of mAbs by single B cell sorting

Cryopreserved human PBMCs were thawed and stained with Live/DEAD Fixable Violet Dead Cell Stain (Life Technologies) for single B cell sorting. After washing, cells were stained with a cocktail of anti-human antibodies, including CD3 (clone SP34-2, BD Biosciences), CD4 (clone OKT4, BioLegend), CD8 (clone RPA-T8, BioLegend), CD14 (clone M5E2, BioLegend), CD20 (clone 2H7, BioLegend), and IgG (G18-145, BD Biosciences), and subsequently stained with fluorescently labeled HCoV-HKU1 S-2P (APC), SARS-CoV S-2P (BV786), and MERS-CoV S-2P (PE) probes (**Figure 1B**). Probe-positive single B cells were sorted into 96-well plates containing lysis solution as previously described^66–68^. Ig heavy-and light-chain mRNAs were reverse-transcribed and amplified by nested PCR using published primers^69^. Paired heavy- and light-chain cDNA sequences were cloned into expression vectors containing constant regions of human Ig heavy (g) and light (k or l) chains. IgG was expressed by co-transfecting Expi293 cells with equal amounts of paired heavy and light chain plasmids, followed by antibody purification using Protein A Fast Flow (GE Healthcare) according to the manufacturer’s instructions. Antibody heavy- and light-chain sequences were compared with human Ig germline sequences using IMGT/V-QUEST^70,71^.

### Enzyme-linked immunosorbent assay (ELISA)

96-well Nunc MaxiSorp plates (Thermo Fisher Scientific) were coated with recombinant S-2P (HCoV-HKU1, HCoV-OC43, SARS-CoV, or MERS-CoV), HCoV-HKU1 S1, or HCoV-HKU1 RBD proteins at 1 μg/mL in 1× PBS at 4°C overnight. Following a 90-min-blocking step with PBS-Tween + 5% non-fat dairy milk, plates were incubated with either serial dilutions of mAbs or heat-inactivated human serum samples (1:200, 4-fold, 8x) for 1 h. Anti-human IgG horseradish peroxidase (HRP)-conjugated secondary antibodies (Jackson Laboratory) were used. To detect binding IgG, plates were developed with 3,3’,5,5’-tetramethylbenzidine (TMB) (KPL) and stopped with 1 N sulfuric acid. OD450 values were collected using a SpectraMax Paradigm (Molecular Device). Endpoint titers were calculated at 4× background of secondary-alone wells.

### Competition binding analysis by ELISA

mAb antibody mapping was performed using competition ELISA, using previously published methods^66,72^. Briefly, mAbs were biotinylated using the EZ-Link Sulfo-NHS-Biotinylation Kit (Thermo Fisher Scientific) and titrated on HCoV-HKU1 S-2P-coated plates. Avidin D-HRP conjugate (Vector Laboratories) and TMB (KPL) were used for color development. Absorbance at 450 nm (OD450) was measured using a SpectraMax Plus plate reader (Molecular Devices). The concentration of biotinylated mAb within the linear range of the titration curve was chosen for the competition ELISA. Unlabeled competitor mAbs were serially diluted and added to S-2P-coated plates. After a 30-min incubation at room temperature, biotinylated mAbs were added, and OD readings were recorded. Biotinylated mAb alone served as a binding control. Percent inhibition of binding was calculated as follows: % competition = [1 – (OD with competing mAb)/(OD without competing mAb)] ×100.

### Competition binding analysis by biolayer interferometry (BLI)

mAb epitope mapping was also performed using biolayer interferometry, as previously described^73^, on the Octet HTX instrument (ForteBio). Briefly, His-tagged HCoV-HKU1 S-2P was loaded onto hydrated Anti-Penta-HIS (HIS1K) biosensors (ForteBio) for 5 min. The sensors were then incubated in blocking buffer (1% BSA, 0.01% Tween-20, 0.02% NaN_3_ in PBS) for 1 min prior to incubation with competitor mAbs for 5 min. After binding of the competitor mAbs, sensors were incubated in blocking buffer for 1 min, followed by incubation with analyte mAbs for 5 min. Percent competition was calculated using the following equation: % competition = [1 – (Analyte binding in the presence of competitor mAb)/(Analyte binding in the absence of competitor mAb)] × 100.

### mAb affinity measurements by BLI

Anti-hIgG Fc capture sensortips (FortéBio) were soaked in running buffer composed of 10 mM HEPES pH 7.5, 150 mM NaCl, 3 mM EDTA, 0.005% Tween 20, and 1 mg/mL BSA for 20 min before being used to capture mAbs H501-003, -007, -008, -009, -010, -012, -013, -014, -015, -016, -018, -020, or -022 to a level of ∼1.4 nm in an Octet RED96 (FortéBio). Tips were then dipped in serial dilutions of either 50–1.5625 nM HCoV-HKU1 NTD (for H501-007 and -010), 10-0.3125 nM HCoV-HKU1 RBD (for H501-003, -008, -009, -012, -013, -014, -015, -016, H501-018, -020, and -022), or 200–3.125 nM HCoV-HKU1 S-2P (for H501-101) before being dipped into wells containing only running buffer to measure dissociation. Data were reference-subtracted in Octet Data Analysis software v11.1 (FortéBio) and fit to a 1:1 binding model.

### mAb affinity measurements by surface plasmon resonance (SPR)

Affinity-purified Fc- and His-tagged HCoV-HKU1 RBD or HCoV-OC43 S1-CTD was immobilized to a single flow cell of an NTA sensor chip at a level of ∼300 response units (RUs) per cycle using a Biacore X100 (GE Healthcare). The sensor chip was doubly regenerated using 0.35 M EDTA and 0.1 M NaOH followed by regeneration with 0.5 mM NiCl2. Three samples containing only HBS-P pH 8.0 running buffer were injected over both ligand and reference flow cells, followed by H501-008 Fab serially diluted from 25–0.78125 nM, with a replicate of the 6.25 nM concentration. The resulting data were double-reference-subtracted and fit to a 1:1 binding model using the Biacore X100 Evaluation software. Twin-Strep-tagged and His-tagged HCoV-HKU1 S or HCoV-OC43 S was immobilized to a single flow cell of an NTA sensor chip at a level of ∼250 RUs per cycle, and H501-101 Fab binding was measured over a serial dilution from 25– 0.78125 nM with a replicate of the 6.25 nM concentration. Sensor chip regeneration, reference subtraction, and data processing were performed under the same conditions described above.

### HCoV-HKU1 pseudovirus neutralization assay

Pseudotyped HCoV-HKU1 coronavirus was generated as described elsewhere with minor modifications^39,74^. Briefly, the full-length HCoV-HKU1 S gene (GenBank ID: DQ339101) or HCoV-HKU1 S with indicated amino acid substitutions lacking the native signal peptide and endoplasmic retention signal sequence was cloned into the pcDNA3.1 expression vector with a tissue plasminogen activator signal sequence (MDAMKRGLCCVLLLCGAVFVSA). A total of 10 μg of plasmid was transfected into HEK293T cells using PEI (Polysciences) at a 1:3 ratio in OPTI-Minimum Essential Medium (Opti-MEM, Gibco, Thermo Fisher Scientific). At 24 h post-transfection, cells were infected with VSVΔG/Luc virus and incubated for 2 h at 37°C with 5% CO_2_. The cells were washed thrice with PBS, and complete DMEM (DMEM with 10% FBS and 1% penicillin-streptomycin) containing anti-VSV-G antibody (I1 mouse hybridoma supernatant diluted 1:25 from ATCC-CRL-2700) was added to the cells to reduce parental virus background and incubated for 24 h. Cell supernatants were harvested 24 h post-infection, cell debris was removed by centrifugation at 3,000 rpm for 10 min, filtered (0.45 μm), and aliquots were stored at -80°C until used. For pseudovirus titration, HEK293-TMPRSS2 cells grown in 96-well black-and-white plates were infected with 50 μL of serially diluted virus stock (2-fold dilutions). After 24 h incubation at 37°C in a CO_2_ incubator, cells were lysed using cell lysis buffer (Promega) following the manufacturer’s instructions, and subsequently, 50 μL of luciferase substrate (Promega) was added to the cells. Relative light unit (RLU) values were calculated and plotted against log10 virus dilutions in GraphPad Prism.

For pseudovirus neutralization assays, HEK293T cells at 80% confluency in 100mm plates were transfected with 10 μg of TMPRSS2 plasmid and incubated for 6 h. Transfected cells (HEK293T-TMPRSS2) were trypsinized, counted, and reseeded at ∼20,000 cells/well in 96-well black and white plates (Revvity) and incubated at 37°C with 5% CO_2_ overnight. The following day, mAb serial dilutions (4-fold, 8×, starting at 20 μg/mL) were prepared in DMEM (Gibco, ThermoFisher Scientific) in a final volume of 90 μL. A total of 90 μL of HCoV-HKU1 pseudovirus (100,000 RLU) was added to each well, and the pseudovirus/Ab mixture was incubated at 37°C for 45 min. Then, 50 μL of pseudovirus/Ab mixture was added to HEK293T-TMPRSS2 cells and incubated for 2 h at 37°C, followed by the addition of 100 μL of DMEM with 10% FBS and 1% penicillin-streptomycin. At 20 h post-infection, the cells were lysed using cell lysis buffer (Promega) following the manufacturer’s instructions, and subsequently, 50 μL of luciferase substrate (Promega) was added to the cells. Luciferase readout (RLU) was measured using a SpectraMax iD5 plate reader (Molecular Devices). RLU values in triplicate were plotted and normalized in GraphPad Prism, with a baseline value of pseudovirus alone set to 0% neutralization and uninfected cells representing 100% neutralization. Nonlinear regression was employed to calculate the IC_50_ values from the fitted curves.

### HCoV-HKU1 live virus assay

HCoV-HKU1-B was recovered from a patient’s nasopharyngeal material and subsequently propagated in normal human bronchial epithelial (NHBE) cells (Lonza, CC-2540). For neutralization studies, 50 µL of virus was mixed with serial concentrations (10 µg/mL, 10-fold, 4X) of mAbs and incubated at 34°C for 30 min. The virus-antibody mixtures were then applied to NHBE cells and allowed to adsorb for 2 h at 34°C. After adsorption, the inoculum was removed, cells were washed twice with phosphate-buffered saline (PBS), and cultures were maintained at 34°C. Apical supernatants were collected in 200 µL PBS at 24, 48, 72, and 96 h post-infection (hpi). At 96 hpi, cell lysates were harvested for analysis of viral nucleocapsid (N) protein expression by Western blotting. Viral RNA was isolated from the apical washes using the Quick-RNA Viral Kit (Zymo Research) and eluted in 20 µL nuclease-free water. Quantification was performed using 2 µL of each RNA preparation. A standard curve was generated using freshly synthesized in vitro–transcribed RNA corresponding to the HKU1 N gene. Neutralizing efficacy of the mAbs was assessed by determining reductions in viral RNA levels over time and across antibody concentrations. Western blot analysis was carried out using anti-HKU1 N antibody (Sino Biological, 40642-T62) and anti-GAPDH antibody (GeneTex, GTX627408) as a loading control.

### HCoV-OC43 neutralization assay

Huh7 cells were seeded at 20,000 cells/well in a Poly-L-Lysine (Gibco, Thermo Fisher Scientific)-coated 96-well plate. The following day, mAbs (H501-008, H501-101, and CDC2-C2^75^) were serially diluted (4-fold, 9× starting at 40 µg/mL) in Opti-MEM (Gibco, Thermo Fisher Scientific) containing 2% FBS, NEAA (Gibco), and anti/anti (Thermo Fisher Scientific). Equal volumes of HCoV-OC43 (VR-1558 strain) (MOI = 0.025) were added to each well and incubated at 37°C for 1 h. Following incubation, 50 µl of each virus/antibody series was added to Huh-7 cells in quadruplicates and incubated at 32°C for 1 h. After 1 h, cells were overlaid with Optimem containing 1% carboxymethyl cellulose, 2% FBS, NEAA, and anti/anti. After 72 h incubation at 32°C, cells were fixed with 10% buffered formalin (Thermo Fisher Scientific), permeabilized using 0.1% Triton X-100 (Sigma-Aldrich), and blocked in PBS containing 1% BSA and 0.2% skim milk. Endogenous peroxidase activity was quenched with 3% hydrogen peroxide (Thermo Fisher Scientific). HCoV-OC43 antigen was detected using a mouse anti-OC43 nucleoprotein primary antibody (Millipore MAB9013) followed by an HRP-conjugated goat anti-mouse secondary antibody (KPL 474-1806) and visualized with DAB reagent (Thermo Fisher Scientific). Infected cell foci were imaged and quantified using a CTL ImmunoSpot ELISpot reader. The effective concentration resulting in a 50% reduction in viral replication was defined as EC_50_. EC_50_ values were calculated from a dose-response curve using a 4-parameter (variable slope) equation (Equation 1 in GraphPad Prism: Y = 100/(1 + 10((LogEC_50_ − X) × HillSlope)), where Y represents % inhibition and X represents antibody concentration. Mock-infected cells were used as a control for 100% inhibition, and cells treated with virus alone were used as the baseline for 0% inhibition.

### OC43-NV in vivo mouse model challenge

6-8 weeks old C57BL/6 mice (Jackson Laboratory) were intranasally challenged with 50 µL of OC43-NV stock (1 × 10^10^ genome copies), administered as 25 µL per nostril. Twelve hours post-challenge, the mice were intraperitoneally administered either H501-008, -101, or -022 at 20mg/kg or PBS. All mice were monitored daily for weight loss, clinical severity, and survival over the course of 2 weeks. Disease severity was measured using clinical scores ranging from 0 to 3, as described previously ^9^. Major changes included body posture, appearance of fur, eye secretions or closure, neurological stimuli reactivity or distress, and animal activity. A score of 3 indicated severe disease, characterized by severe hunching, complete lack of responsiveness and movement even upon stimulation, fully closed eyes, cessation of eating and drinking, severe weight loss, and the presence of severe neurological symptoms. A score of 2 represented moderate disease, including a moderately hunched posture, extensive fur ruffling along the back, decreased food intake, reduced responsiveness to auditory or tactile stimuli, partially closed eyes, possible ocular discharge, lethargy, and persistent labored breathing. A score of 1 was assigned to mild disease, characterized by slight hunching, mildly ruffled fur, and delayed responses to auditory or tactile stimulation while remaining active. Healthy animals exhibiting normal activity and a smooth, shiny coat were assigned a score of 0.

### Cell surface mAb binding assay

HEK293T cells were transiently transfected with plasmids encoding full-length S proteins from HCoV-HKU1, HCoV-OC43, MERS-CoV, SARS-CoV, MHV-CoV, or HCoV-HKU1 with indicated amino acid substitutions, using Lipofectamine 3000 (Thermo Fisher Scientific) according to the manufacturer’s protocol. After 40 h, cells were harvested and incubated with mAbs (1 μg/mL) for 30 min. Following antibody incubation, cells were washed and stained with an allophycocyanin-conjugated anti-human IgG secondary antibody (Jackson ImmunoResearch Laboratories, 709-136-149) for an additional 30 min. Cells were then washed and fixed with 1% paraformaldehyde (Electron Microscopy Sciences). Samples were acquired on a BD LSR Fortessa X-50 flow cytometer (BD Biosciences) and analyzed using FlowJo v10.10.0 (BD Biosciences).

### S-TMPRSS2 binding assay

To determine whether HCoV-HKU1 mAbs interfere with S binding to TMPRSS2, we performed an S-binding assay on HEK293T-TMPRSS2 cells as described elsewhere^76^. Briefly, TMPRSS2 was transiently expressed in HEK293T cells seeded on coverslips (Electron Microscopy Sciences, 72290-02) for immunofluorescence staining or 6-well plates (Costar, 3516) for fluorescence activated cell sorting (FACS). Before flow cytometry analysis, HEK293T-TMPRSS2 cells seeded in 6-well plates were trypsinized and added to round-bottom plates (Costar, 3788). The transfected cells were blocked with 1% BSA for 30 min at 4°C followed by a PBS (Gibco, Thermo Fisher Scientific) wash. Prior to addition of HCoV-HKU1 RBD to TMPRSS2-expressing cells, recombinant protein was incubated independently with 10 μg/mL of HCoV-HKU1 mAbs for 30 min at 4°C. The mAb/CTD mixture was allowed to bind to HEK293T-TMPRSS2 cells for 1 h at 4°C, followed by two PBS washes. For immunofluorescence staining of HCoV-HKU1 RBD, binding was detected using anti-human Alexa Fluor 594 (Invitrogen, A11014), whereas for flow cytometry analysis, staining was performed with anti-human FITC (Bethyl, A80-119F). Following secondary antibody staining, cells were washed with PBS, fixed in 4% PFA and processed independently for confocal imaging or FACS. Images were acquired on a Nikon Eclipse T2 confocal microscope and processed using Nikon Elements v5.21.0, whereas FACS samples were acquired on an iQue 3 (Sartorius) and analyzed using FlowJo v10.10.0.

### X-ray crystallography

HCoV-HKU1 RBD (residues 310–606 of isolate N5) was treated with HRV3C protease, as described above, and the digested protein was passed over Ni-NTA resin to remove cleaved tags and excess protease. Untagged HCoV-HKU1 RBD was then mixed with a molar excess of H501-008 Fab and passed over a Superdex 75 column using 2 mM Tris pH 8.0, 200 mM NaCl, 0.02% NaN_3_. The purified complex was then concentrated to 6.89 mg/mL and used to prepare sitting-drop vapor-diffusion crystallization trays. Crystals grown in 0.1 M Tris pH 8.5, 0.2 M MgCl_2_, and 25% PEG 3350 were soaked in mother liquor supplemented with 20% glycerol and frozen in liquid nitrogen. Diffraction data were collected to a resolution of 2.60 Å at the ALS beamline 5.0.1. Diffraction data were indexed and integrated using iMOSFLM^77^ before being scaled in AIMLESS^78^. The HCoV-HKU1 RBD + H501-008 Fab dataset was phased by molecular replacement in Phaser-MR^79^ using coordinates from PDBs 4YDJ, 3TV3, and 5KWB as search ensembles. The resulting molecular replacement solution was iteratively rebuilt and refined using Coot^80^, ISOLDE^81^, and Phenix^82^. Crystallographic software packages were curated by SBGrid^83^.

### Cryo-EM sample preparation

Fabs were immediately complexed with 6–7 mg/mL HCoV-HKU1 S-2P at a 1.1:1 molar ratio of Fab to S protomer followed by the addition of fluorinated octyl-maltoside (FOM, Anatrace) to a final concentration of 0.02% w/v. Complexes (3 μL) were applied to Quantifoil Cu 1.2/1.3 300 mesh grids that were freshly glow-discharged at 0.38 mbar, 15 mA, for 60 s using a PELCO easiGLOW (Ted Pella). Grids were blotted using a Vitrobot cryo-plunger (Thermo Fisher Scientific) for 3–4 s at 6°C and 100% humidity prior to plunging into liquid ethane.

### Cryo-EM data collection and processing

For HCoV-HKU1 S-2P with H501-008 Fab complex, single-particle data were collected on a Glacios 2 Cryo-TEM equipped with a Falcon 4i detector (Thermo Fisher Scientific). The microscope was operated at 200 kV, 150,000× magnification, and images were collected as EER frames at 0.923 Å/pixel (unbinned) using EPU automated data collection software. Images were collected in counting mode, one shot per hole, applying beam shift with a total exposure of 40 el/Å^2^, and defocus values ranged from 0.7–2.5 μm. For the HCoV-HKU1 S-2P with H501-018 Fab and HCoV-HKU1 S-2P with H501-022 Fab complexes, single-particle data were collected on a Titan Krios G2 Cryo-TEM (Thermo Fisher Scientific). The microscope was operated at 300 kV, 130,000× magnification, and images were collected as EER frames at 0.92 Å/pixel (unbinned) using EPU automated data collection. Images were collected in counting mode, three shots per hole, applying beam shift with a total exposure of 30 el/Å^2^ and defocus values ranging from 0.4–2.5 μm. Additional data collection parameters can be found in Supplementary Table 2.

For all cryo-EM datasets, patch motion correction, patch CTF estimation, and reference-free particle picking using circular picks ranging from 100–300 Å were carried out in cryoSPARC v4.6^35,84^. Particles were extracted using a box size of 480 pixels (HCoV-HKU1 S-2P with H501-018 or H501-022) or 500 pixels (HCoV-HKU1 S-2P with H501-008) and downsampled 4× prior to 2D classification. Particles selected from 2D classification were input for antibody-initio generation assigning 2–4 classes, and heterogeneous refinement to obtain a starting cryo-EM map of S-Fab complexes for downstream homogeneous refinements, local refinements, and global non-uniform refinements. HCoV-HKU1 S-2P with H501-022 Fab was further 3D classified into 4 classes in Relion v5 while applying Blush Regularization^85,86^. To separate out the different states of HCoV-HKU1 S-2P with H501-018 Fab, particles were taken into Relion v5, and local masked 3D classification with Blush Regularization was performed, assigning 3 classes^85,86^. The circular mask was generated from apoferritin (EMD-20026, low-pass filtered to 15 Å, extending by 3 pixels with a soft-edge of 6 pixels) resampled onto one protomer of the S-Fab density to sort out particles with Fabs bound to either 2-up 1-down RBD or 2-down 1-up RBD^87^. Particles classified in Relion were imported back into cryoSPARC for another round of non-uniform refinement. Resolutions of all maps were generated applying the Fourier shell correlation cutoff criterion of 0.143 in cryoSPARC. Data processing statistics and workflow summaries can be found in **Table S2 and Figure S6**, respectively.

### Cryo-EM structure modeling, refinement, and analyses

Coordinates for initial models were fit into cryo-EM map densities using ChimeraX prior to real space refinement and validation^88–90^. PDB codes used for modeling HCoV-HKU1 S were 8Y8A, 8Y8H, and 8Y8G for S with 3 RBD up, 2-up-1-down RBD, and 2-down-1-up RBD, respectively^40^ . For S structures, rigid-body docking was performed in ChimeraX prior to validation in Phenix^88–91^. *N*-linked glycans were modeled where density for glycans was observed near N-X-T/S S sequences. Proline residue substitutions at positions 902, 980, and 1,023 were mutated back to the original sequences followed by real-space refinement in Coot^92^. Starting models of variable heavy (V_H_) and variable light (V_L_) chains were obtained from the IMGT 3D structure database^93^. PDB codes 7T3M and 5O7P^94,95^, respectively, were docked into the V_H_ or V_L_ density of the locally refined map for H-022 Fab bound to S 2P. PDB codes 4FQK and 4QHK^96,97^, respectively, were docked into the locally refined map for H-018 Fab bound to S-2P. Model building and real-space refinement for Fab-RBD structures were carried out in Coot and Phenix ^91,92^. BSA analyses were carried out using PDBePISA^36^. Total BSA was determined by taking the sum of all residue BSA contributions. Interacting residues were identified using a 4 Å search cutoff in Coot and ChimeraX^88–90,92^.

### mAb footprint analysis

HCoV-HKU1 S sequences from genotypes A, B, and C were retrieved from the National Center for Biotechnology Information GenBank database, and the spike sequence of HCoV-HKU1-B used for the live virus assay was obtained by whole-genome sequencing. Multiple sequence alignment was performed in MEGA11^98^ using the MUSCLE algorithm. Aligned sequences were trimmed to represent the RBD region (residues 441–564), and mAb binding residues were highlighted in red (H501-008), purple (H501-018), and green (H501-022).

## Data Availability

Source data supporting the findings of this study are available within this paper and its supplementary files. Source data have also been deposited to Figshare and can be accessed here: 10.6084/m9.figshare.30468065. Additional source data inquiries should be directed to the corresponding author, Kizzmekia Corbett-Helaire. The atomic models and cryo-EM maps generated for H501-018 and -022 bound to HCoV-HKU1 S-2P have been deposited in the Protein Data Bank (PDB; http://www.rcsb.org/) and the Electron Microscopy Data Bank (EMDB; http://www.emdataresource.org/) under accession codes PDB: 9YGN, 9YGO, 9YGP, 9YGQ, and 9YGR and EMD-72935, EMD-72936, EMD-72937, EMD-72938, and EMD-72939, respectively. Additionally, the atomic coordinates for H501-008 bound to RBD are deposited in the Protein Data Bank under accession code 9YXW, and the cryo-EM map for H501-008 bound to HCoV-HKU1 S-2P is deposited in the Electron Microscopy Data Bank under accession code EMD-72934.

## Material and Resource Availability

Requests for reagents and resources must be directed to the corresponding author, Kizzmekia Corbett-Helaire, and will be provided upon execution of a Material Transfer Agreement. If the material was obtained under use restriction, the inquiry will be forwarded to the appropriate party.

## Acknowledgments

We thank Kaitlyn Morabito for project management, Fidan Baycora and Monique Young for administrative support, Emily Hobbs for grant support, and Harvard and HHMI’s Office of Technology Development for materials transfer and collaboration agreement support. This work was supported, in part, by a Chan Zuckerberg Initiative Science Diversity Leadership Grant (No. 2022-310965 to KSC), Howard Hughes Medical Institute Freeman Hrabowski Scholars grants (to KSC and COB), start-up funds from Harvard T.H. Chan School of Public Health (to KSC), in-kind gifts of laboratory equipment, consumables, and supplies from Corning, Inc. (to KSC), the Howard Hughes Medical Institute Emerging Pathogens Initiative (to COB), and the VRC, an intramural division of NIAID, NIH. KSC’s research fellowship was partially funded by the Undergraduate Scholarship Program, Office of Intramural Training and Education, Office of the Director, NIH. Additionally, COB is supported by the Rita Allen Foundation and the Pew Charitable Trusts Biomedical Scholars Program and is a Chan Zuckerberg Biohub investigator. BLI and SPR studies to assess mAb binding affinity and X-ray crystallographic studies were supported in part by NIAID (R01-AI127521 to JSM). HCoV-OC43 neutralization studies were funded by NIAID (R01 AI132178 to RB and TPS). Cryo-EM structural studies of H501-008 were supported by NIAID (R01-AI127521 to ABW). Propagation of OC43-NV isolate for challenge studies was supported by AAI Restore from the American Association of Immunologists (to PPM). HCoV-HKU1 live virus neutralization studies were supported by start-up funds from Boston University (to MS).

## Author Contributions

HY, BSG, and KSC conceptualized the study. HY and BSG spearheaded acquisition of the clinical sample. LW, WS, WK, and YZ isolated and performed binding characterization of the mAbs, under the direction of TP and JM. JJ optimized and completed the HCoV-HKU1 pseudovirus assays and mAb surface staining and TMPRSS2 binding competition analyses with technical assistance from CH and SM. JJ and SM performed the HCoV-OC43 challenge experiment with assistance from TD and PPM. CMS and JJ completed the HCoV-HKU1 live virus assays/analysis under the direction of MS. SV completed cryo-EM studies of all mAbs bound to S-2P with technical assistance from MRE, under the direction of COB. DW performed BLI, SPR, and X-ray crystallographic analyses under the direction of JSM. TPS and AJB completed HCoV-OC43 live-virus neutralization assays, under the direction of RB. OA and OR completed serum-binding analyses. CKOD performed multiple sequence alignment of HCoV-HKU1 S and generated the antibody binding footprint. RNK completed cryo-EM studies of H501-008 bound to HCoV-HKU1 RBD under the direction of ABW. KSC supervised and administered the work. JJ outlined and wrote the original manuscript with significant input from LW, SV, and KSC. JJ and KSC revised and polished the manuscript with input from all co-authors.

