## Supplementary Figures for "Structural Definition of Neutralizing Epitopes within and around the HKU1 Receptor-binding Site Reveals Cell Context–Dependent Neutralization"

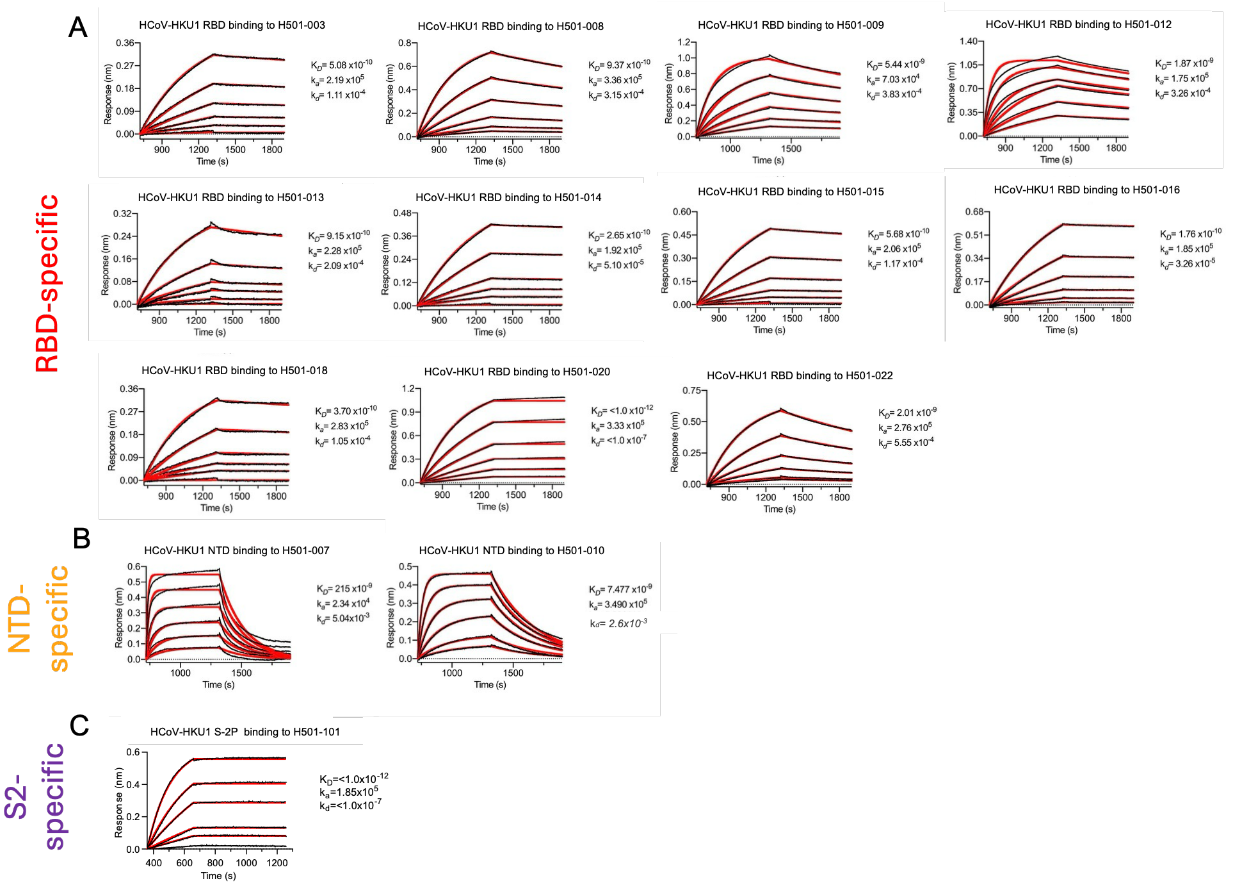


**Figure S1. Binding affinity of HCoV-HKU1 mAbs.** Binding affinities of RBD- **(A)**, NTD- **(B)**, and S2-specific **(C)** mAbs for HCoV-HKU1 RBD, NTD, and S-2P, respectively. mAbs were immobilized on biosensors and analyzed using a 1:1 binding kinetic model by biolayer interferometry. $K_D$ (M), $k_a$ ($M^{-1}s^{-1}$), and $k_d$ ($s^{-1}$) values were calculated (**Table 1**).

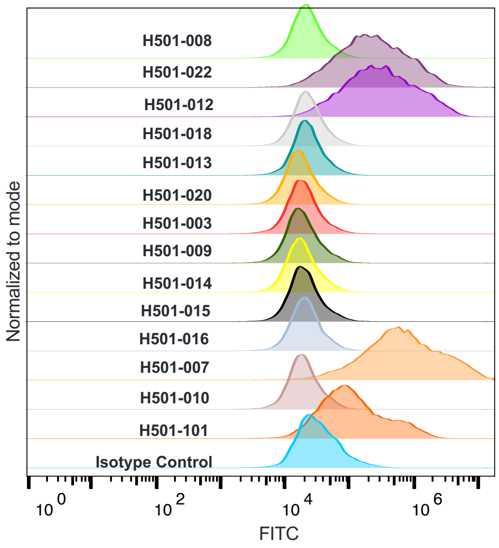


**Figure S2. Binding of HCoV-HKU1 mAbs to HCoV-HKU1 genotype A.** HCoV-HKU1 genotype A S protein expressed on the surface of HEK293T cells was stained with HCoV-HKU1 mAbs, followed by detection using a FITC-conjugated secondary antibody. Histograms show FITC fluorescence intensity for each mAb. An isotype control is shown at the bottom.

| Ab1 | Ab2 | | | | | | | | | | | | |
|---|---|---|---|---|---|---|---|---|---|---|---|---|---|
| | H501-008 | H501-022 | H501-012 | H501-018 | H501-013 | H501-020 | H501-003 | H501-009 | H501-014 | H501-015 | H501-016 | H501-007 | H501-010 |
| H501-008 | 94 | -9 | -17 | 22 | 19 | 13 | -11 | 2 | 0 | 7 | 3 | 15 | -10 |
| H501-022 | 18 | 89 | 95 | 34 | 23 | 11 | 1 | 18 | 19 | 14 | 8 | 18 | 0 |
| H501-012 | 6 | 88 | 90 | 18 | 27 | 22 | 29 | 28 | 17 | 18 | 24 | 21 | -3 |
| H501-018 | 38 | 49 | 49 | 94 | 96 | 92 | 105 | 95 | 97 | 95 | 91 | 19 | 11 |
| H501-013 | 33 | 46 | 38 | 96 | 97 | 97 | 100 | 101 | 100 | 104 | 91 | 11 | 15 |
| H501-020 | 28 | 43 | 44 | 97 | 98 | 97 | 101 | 97 | 98 | 102 | 92 | 13 | -4 |
| H501-003 | 27 | 29 | 26 | 82 | 82 | 77 | 83 | 84 | 80 | 78 | 77 | 9 | -13 |
| H501-009 | 16 | 16 | 27 | 93 | 96 | 94 | 103 | 95 | 96 | 92 | 89 | -1 | -11 |
| H501-014 | 24 | 34 | 24 | 95 | 88 | 88 | 91 | 92 | 90 | 87 | 79 | 7 | -7 |
| H501-015 | 30 | 46 | 46 | 93 | 98 | 95 | 110 | 99 | 100 | 93 | 104 | 18 | -1 |
| H501-016 | 11 | 29 | 36 | 96 | 99 | 95 | 135 | 101 | 99 | 103 | 92 | 8 | -7 |
| H501-007 | 43 | 28 | 14 | 27 | 33 | 26 | 36 | 33 | 26 | 29 | 19 | 93 | 42 |
| H501-010 | 24 | 32 | 26 | 30 | 35 | 35 | 42 | 25 | 18 | 27 | 16 | 43 | 100 |

**% Inhibition** >90% | 80-89.99% | 60-79.99% | 50-59.99% | <50%

**Figure S3. HCoV-HKU1 mAb competition map**. HCoV-HKU1 RBD- (red text labels) and NTD-specific (orange text labels) mAbs were assessed for competition binding to HCoV-HKU1 S-2P in the presence of mAbs listed in the leftmost column. Analyses were completed by biolayer interferometry. Percent inhibition of analyte mAb binding by competitor mAb is indicated by color. Dark red = >90% competition; red = 80–99.99% competition; yellow = 60–79.99% competition; uncolored = <50% competition.

**Table S1**. mAb Heavy and light gene sequences

| mAbs | VH | % identity | CDRH3 length | CDRH3 | VL | % identity | CDRL3 length | CDRL3 |
|---|---|---|---|---|---|---|---|---|
| **H501-003** | IGHV4-59 | 95.09 | 19 | CASAPVLGSSGPKGYYGLEVW | IGLV1-47 | 98.25 | 11 | CAAWDDSLSGYVF |
| **H501-008** | IGHV3-30 | 88.89 | 24 | CAREGPMYYDAWSGRSYYNNYALDVW | IGLV1-51 | 93.33 | 11 | CGTWDSSLSAGVF |
| **H501-009** | IGHV4-61 | 92.98 | 15 | CARSSGQIPKTEAFQHW | IGLV1-47 | 97.54 | 11 | CATWDDSLRTPVF |
| **H501-012** | IGHV2-5 | 97.59 | 15 | CVYSMSRPVAGTVIDHW | IGKV1-39 | 94.62 | 9 | CQQSYSTPPTF |
| **H501-013** | IGHV4-61 | 94.74 | 14 | CARFPGLPKTNAFDIW | IGLV1-47 | 95.09 | 11 | CATWDDSLSCPVF |
| **H501-014** | IGHV4-59 | 93.33 | 14 | CARGGFLKNSDAFDIW | IGLV1-47 | 95.79 | 11 | CAAWDDSLTGPVF |
| **H501-015** | IGHV4-59 | 95.09 | 13 | CARGSWARRLEVDNW | IGLV1-47 | 97.19 | 11 | CATWDDNLGAWVF |
| **H501-016** | IGHV3-48 | 93.06 | 14 | CATHSSNWVYDAFDIW | IGKV1-15 | 93.91 | 12 | CQQYDIWPRLTF |
| **H501-018** | IGHV4-59 | 94.97 | 13 | CARGTWSRRLEVDLW | IGLV1-47 | 98.60 | 11 | CATWDDSLNSWVF |
| **H501-020** | IGHV4-59 | 93.33 | 14 | CARGEWERRLGAFDIW | IGLV1-47 | 94.39 | 11 | CSVWDDSLSAVAF |
| **H501-022** | IGHV2-5 | 96.56 | 15 | CVYSMSRPVAGGVIDHW | IGKV1-39 | 93.19 | 9 | CQQSYSSPPTF |
| **H501-007** | IGHV6-10 | 90.57 | 16 | CARVVVRYGWNDNWFDPW | IGKV1-39 | 91.40 | 9 | CQQSTGTPTF |
| **H501-010** | IGHV5-51 | 96.88 | 17 | CARLRDEGADYYYYGMDVW | IGKV1-9 | 98.92 | 10 | CQQLNSYSLGF |
| **H501-101** | IGHV1-46 | 90.62 | 16 | CARGVDLWGNFYTYFDYW | IGKV3-20 | 90.07 | 9 | CQQHDASPTF |

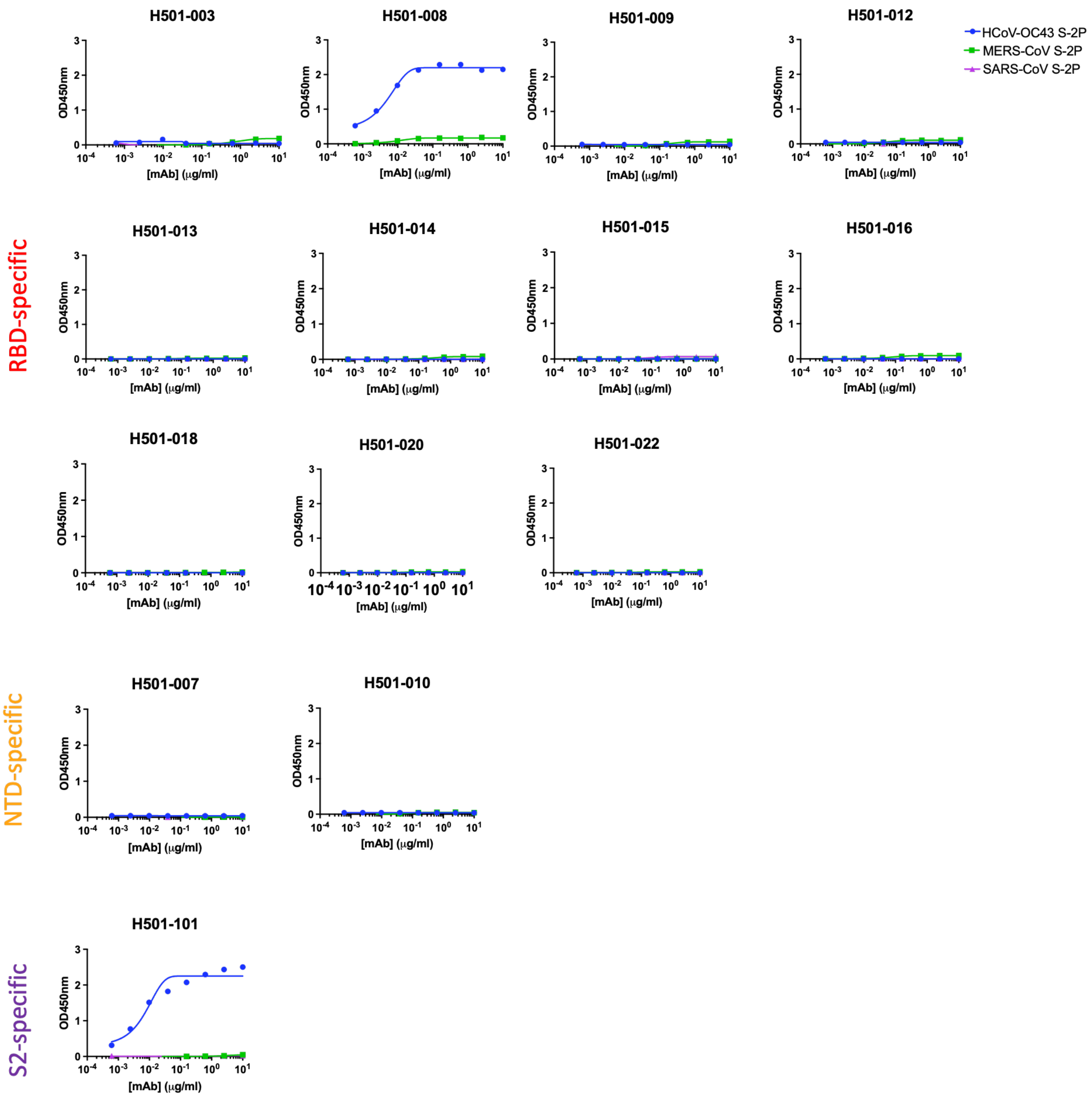


**Figure S4. Binding of HCoV-HKU1 mAbs to beta-CoV S proteins.** Binding of HCoV-HKU1 mAbs to HCoV-OC43 S-2P (blue), MERS-CoV S-2P (green), and SARS-CoV S-2P (purple) was assessed by ELISA. H501-008 and H501-101 bound HCoV-OC43 S-2P, whereas all other mAbs showed no detectable binding to HCoV-OC43, MERS-CoV, or SARS-CoV S-2P. For H501-008 and H501-101, MERS-CoV and SARS-CoV S-2P binding curves remained at baseline, indicating no detectable binding.

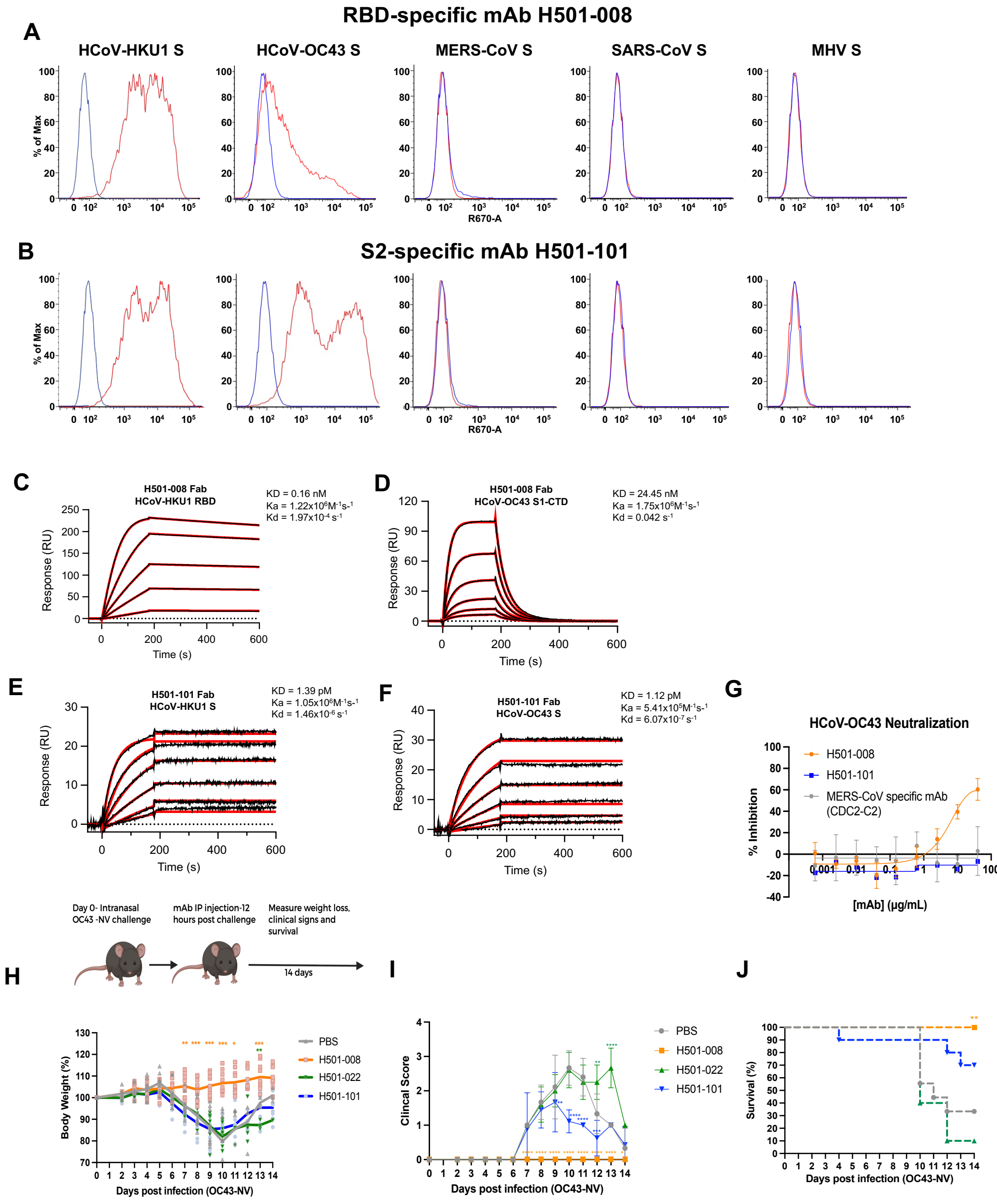


**Figure S5. Human CoV cross-reactivity of RBD-specific and S2-specific mAbs. (A)** RBD-specific mAb H501-008 and **(B)** S2-specific mAb H501-101 were assessed for binding to cell surface-expressed S proteins from HCoV-HKU1, HCoV-OC43, MHV-CoV, MERS-CoV, and SARS-CoV. The percentage of maximum binding to S-transfected cells (red) compared with untransfected cells (blue) is shown. **(C–F)** Binding affinities of H501-008 Fab to **(C)** HCoV-HKU1 RBD and **(D)** HCoV-OC43 S1-CTD, and of H501-101 Fab to **(E)** HCoV-HKU1 S and **(F)** HCoV-OC43 S, were assessed by surface plasmon resonance. **(G)** The ability of H501-008 (orange circles) and H501-101 (blue squares) to neutralize native HCoV-OC43 was assessed. The MERS-

CoV-specific mAb CDC2-C2 (gray circles) was used as a negative control. Error bars represent the standard deviation of three individual measurements. **(H, upper panel)** Experimental outline for assessing whether HCoV-OC43 cross-reactive mAbs protect mice from HCoV-OC43 infection. Mice (n=10) were inoculated intranasally with the OC43 NV strain, followed by intraperitoneal administration of 20 mg/kg HCoV-HKU1 mAbs. Mice were monitored for 14 days and assessed for changes in **(H, lower panel)** body weight, **(I)** clinical score, and **(J)** survival. Body weight and clinical scores were compared using a mixed-effects two-way ANOVA with Dunnett's post-hoc test, with Day 0 body weight normalized to 100% for each mouse. Survival curves were compared using the log-rank Mantel-Cox comparison test. Significant differences compared to PBS control are indicated $^*P \leq 0.05$, $^{**}P \leq 0.01$, $^{***}P \leq 0.001$, $^{****}P \leq 0.0001$. Data are from one experiment with 10 mice per group, and error bars represent SD.

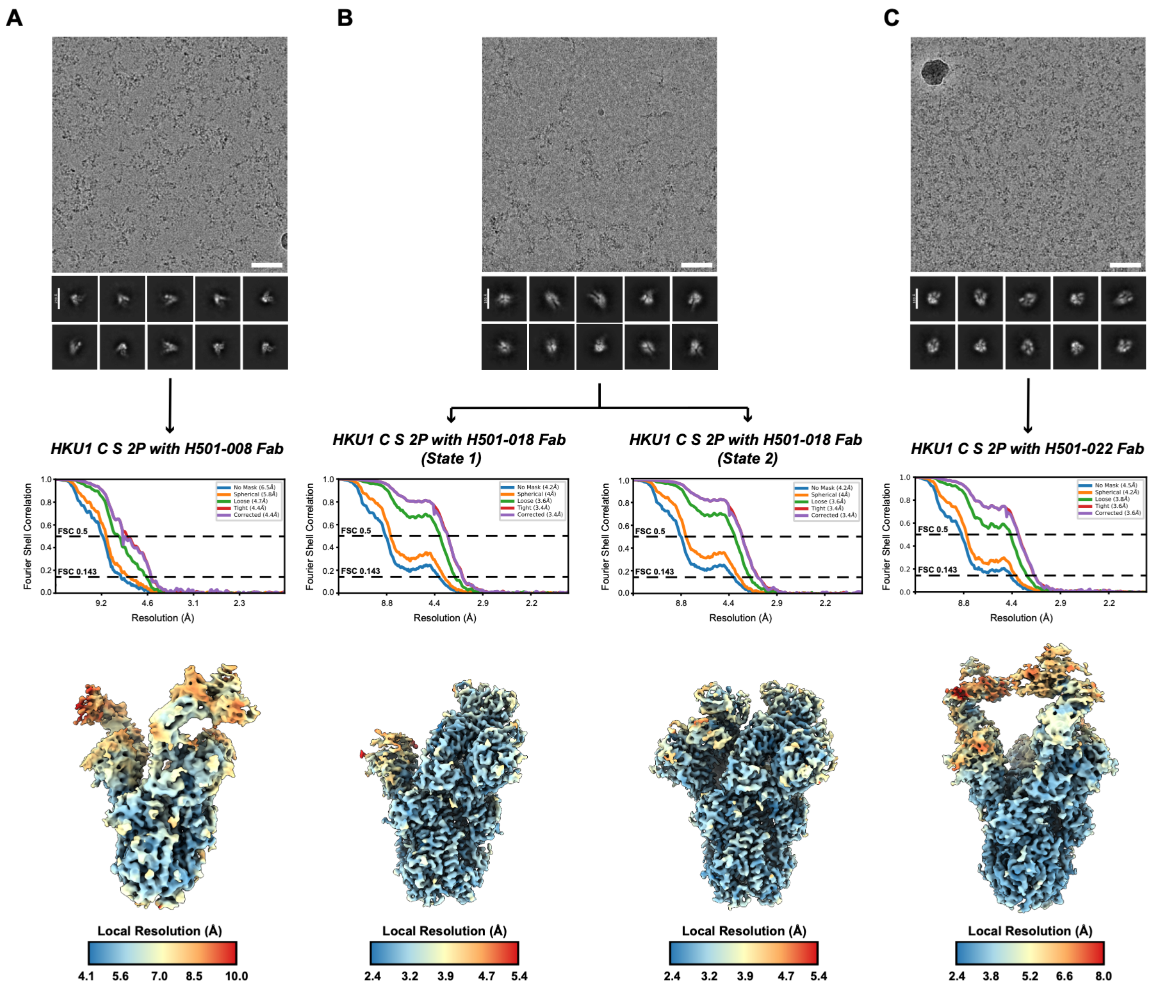


**Figure S6. Cryo-EM data processing summary of H501-008, H501-018, and H501-022 complexes. (a-c)** Representative micrographs, 2D class averages, Fourier Shell Correlation (FSC) plots, and final map reconstructions of H501-008 Fab – S-2P **(A)**, H501-018 Fab – S-2P **(B)**, and H501-022 Fab – S-2P **(C)** complexes. Two states were observed for H501-018 Fab complexed to S-2P **(B)**; one state where there are two S1 RBDs facing up and one RBD down (State 1), and another state with two RBDs facing down and one RBD up (State 2). Scalebars (white) for representative micrographs are 50 nm.

**Table S2. X-ray Crystallography data collection, refinement, and validation statistics**

| | HCoV-HKU1 RBD + H501-008 |
|---|---|
| PDB ID | 9YXW |
| *Data collection* | |
| Space group | $C2_1$ |
| Cell dimensions | |
| a, b, c (Å) | 104.5, 71.8, 174.5 |
| α, β, γ (°) | 90.0, 94.6, 90.0 |
| Resolution (Å) | 59.09-2.60 (2.71-2.60) |
| $I/\sigma I$ | 4.8 (2.2) |
| CC1/2 | 0.919 (0.703) |
| Completeness (%) | 99.8 (99.6) |
| Redundancy | 3.1 (3.3) |
| *Refinement* | |
| $R_{work}/R_{free}$ (%) | 25.2/26.7 |
| No. atoms | |
| Protein | 5480 |
| Ligand | 0 |
| Water | 179 |
| Average B-factors | |
| Protein | 116.32 |
| Solvent | 55.24 |
| R.m.s deviations | |
| Bond lengths (Å) | 0.009 |
| Bond angles (Å) | 1.44 |
| Clashscore | 13.54 |
| Ramachandran | |
| Favored (%) | 96.5 |
| Allowed (%) | 3.4 |
| Outliers (%) | 0.1 |

### Table S3. Cryo-EM data collection, refinement and validation statistics

| | HCoV-HKU1 S-2P H501-008 (EMDB-72934) | HCoV-HKU1 S-2P H501-018 (State 1) (EMDB-72935) (PDB 9YGN) | HCoV-HKU1 S-2P H501-018 (State 2) (EMDB-72936) (PDB 9YGO) | HCoV-HKU1 S-RBD H501-018 (local refine) (EMDB-72937) (PDB 9YGP) | HCoV-HKU1 S-2P H501-022 (EMDB-72938) (PDB 9YGQ) | HCoV-HKU1 S-RBD H501-022 (local refine) (EMDB-72939) (PDB 9YGR) |
|---|---|---|---|---|---|---|
| **Data collection and processing** | | | | | | |
| Magnification | 150000x | 130000x | 130000x | 130000x | 130000x | 130000x |
| Voltage (kV) | 200 | 300 | 300 | 300 | 300 | 300 |
| Electron exposure (e–/Å$^2$) | 40 | 30 | 30 | 30 | 30 | 30 |
| Defocus range (μm) | 0.7 – 2.5 | 0.4 – 2.5 | 0.4 – 2.5 | 0.4 – 2.5 | 0.4 – 2.5 | 0.4 – 2.5 |
| Pixel size (Å) | 0.92 | 0.92 | 0.92 | 0.92 | 0.92 | 0.92 |
| Symmetry imposed | C1 | C1 | C1 | C1 | C1 | C1 |
| Initial particle images (no.) | 860660 | 2019273 | 2019273 | 2019273 | 3417316 | 3417316 |
| Final particle images (no.) | 54647 | 54081 | 54468 | 141143 | 44565 | 139880 |
| Map resolution (Å) FSC threshold 0.143 (unmasked/masked) | 6.5/4.4 | 4.2/3.4 | 4.2/3.4 | 4.3/3.1 | 4.5/3.6 | 6.3/3.9 |
| Map resolution range (Å) | 4.1 – 12.0 | 3.3 – 8.3 | 3.2 – 4.9 | 3.0 – 3.6 | 3.4 – 8.3 | 3.4 – 4.5 |
| **Refinement** | | | | | | |
| Initial model used (PDB code) | | 8Y8H, 4FQK, 4QHK | 8Y8G, 4FQK, 4QHK | 8Y8G, 4FQK, 4QHK | 8Y8I, 7T3M, 5O7P | 8Y8A, 73TM, 5O7P |
| Model resolution (Å) | | | | | | |
| FSC threshold 0.143 | | 3.4 | 3.4 | 3.2 | 3.6 | 3.8 |
| Model resolution range (Å) | | 3.3 - 8.3 | 3.2 – 4.9 | 3.0 – 3.6 | 3.4 – 8.3 | 3.4 – 4.5 |
| Map sharpening *B* factor (Å$^2$) | | 51.2 | 55.0 | 61 | 50.6 | 74.9 |
| Model composition | | | | | | |
| Non-hydrogen atoms | | 32850 | 33115 | 7414 | 31421 | 3312 |
| Protein residues | | 4232 | 4225 | 922 | 4023 | 432 |
| Ligands | | 21 | 24 | 14 | 22 | 0 |
| *B* factors (Å$^2$) | | | | | | |
| Protein | | 101.82 | 75.32 | 79.94 | 112.78 | 67.43 |
| Ligand | | 30.0 | 105.20 | 30.0 | 30.0 | 0 |
| R.m.s. deviations | | | | | | |
| Bond lengths (Å) | | 0.004 | 0.005 | 0.004 | 0.006 | 0.008 |
| Bond angles (°) | | 0.932 | 1.076 | 0.925 | 0.977 | 1.093 |
| Validation | | | | | | |
| MolProbity score | | 1.86 | 1.89 | 1.70 | 2.10 | 1.85 |
| Clashscore | | 7.08 | 8.65 | 4.69 | 8.07 | 7.57 |
| Poor rotamers (%) | | 1.56 | 1.26 | 1.82 | 2.69 | 1.05 |
| Ramachandran plot | | | | | | |
| Favored (%) | | 95.40 | 94.98 | 96.14 | 95.17 | 93.75 |
| Allowed (%) | | 4.60 | 5.02 | 3.86 | 4.83 | 6.25 |
| Disallowed (%) | | 0.00 | 0.00 | 0.00 | 0.00 | 0 |

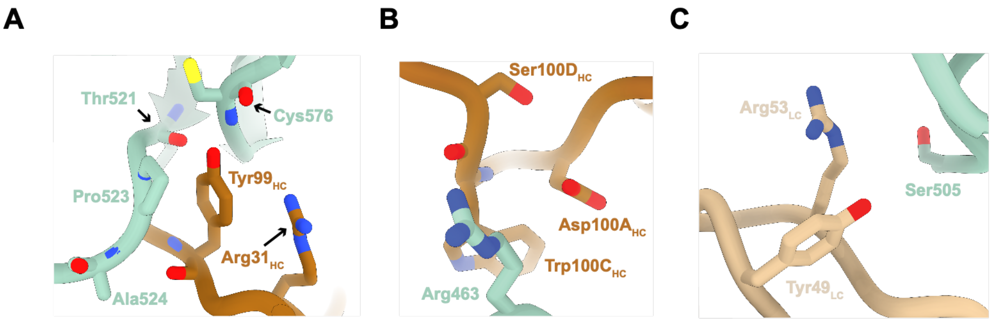


**D**

| | Receptor Binding Domain | | | | | | | | | | | | | | | | | | | | | | | | | | | | |
|---|---|---|---|---|---|---|---|---|---|---|---|---|---|---|---|---|---|---|---|---|---|---|---|---|---|---|---|---|---|
| Residue Number | 442 | 443 | 444 | 445 | 446 | 447 | 448 | 449 | 477 | 478 | 479 | 480 | 481 | 482 | 493 | 494 | 495 | 496 | 497 | 498 | 499 | 500 | 501 | 502 | 563 | 564 | 565 | 566 | 567 |
| HCoV-HKU1 C S (GenBank DQ339101) | S | S | W | N | R | R | Y | G | A | D | P | S | V | V | A | I | C | P | A | G | T | K | Y | R | S | S | C | F | C |
| HCoV-HKU1 A S (GenBank Q5MQD0) | S | S | W | N | R | R | Y | G | A | K | P | S | F | A | A | S | C | P | I | G | T | N | Y | R | V | S | C | L | C |
| HCoV-HKU1 B S (GenBank Q14EB0) | S | S | W | N | R | R | Y | G | A | D | P | S | V | V | A | I | C | P | A | G | T | K | Y | R | S | S | C | S | C |
| HCoV-OC43 S (GenBank AIL49484) | S | T | W | N | K | R | F | G | K | L | N | S | S | - | G | T | C | P | A | G | T | N | Y | L | N | P | C | T | C |
| Buried Surface Area (Å$^2$) | 0 | 20.2 | 0 | 0 | 20.3 | 115 | 0.2 | 0 | 0 | 20.6 | 8.7 | 69 | 9 | 0 | 0 | 53.2 | 18.5 | 41.9 | 93.5 | 36.4 | 0 | 0 | 15.4 | 0 | 36.5 | 45.5 | 6.6 | 0 | 0 |

**Figure S7. Comparisons of HCoV-OC43 S with H501-008 binding residues in HCoV-HKU1 S. (A-C)** Conserved residues from HCoV-OC43 S1 C-terminal domain (S1-CTD) (PDB 7SB3) are modeled near H501-008 by superimposing HCoV-OC43 S1-CTD onto the crystal structure of the HCoV-HKU1 RBD – H501-008 complex. **(D)** Sequence alignment of HCoV-HKU1 S with other HCoV spike sequences. Buried Surface Area (BSA) between HCoV-HKU1 S1-CTD and H501-008 is color-coded based on relative BSA values.

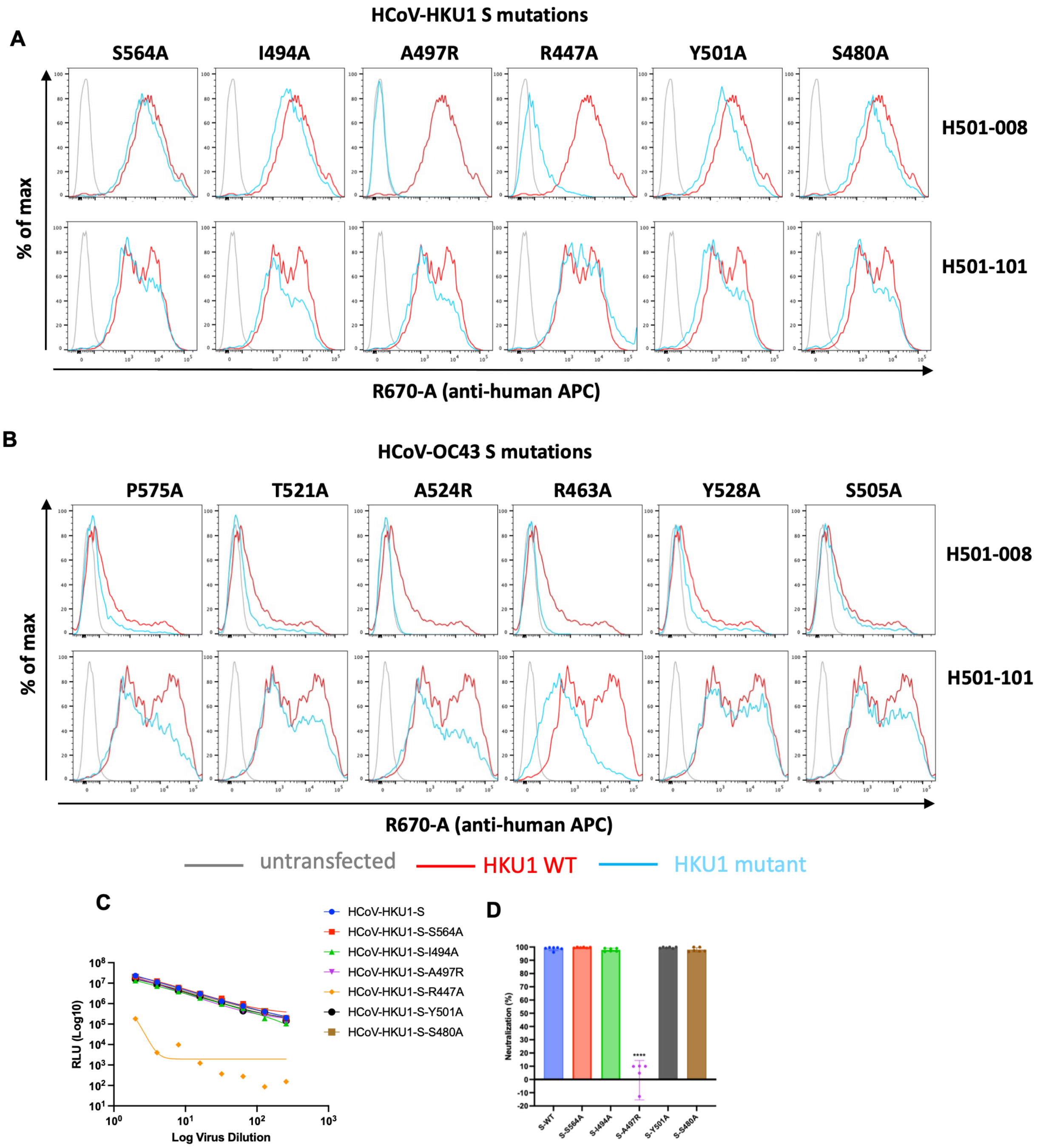


**Figure S8. HCoV-HKU1/OC43-specific mAb H501-008 binding residues. (A-B)** H501-008 and -101 were assessed for binding to cell surface-expressed HCoV-HKU1 **(A)** or HCoV-OC43 **(B)** S with indicated mutations, as revealed by crystal structure in Fig. 2. Percentage of maximum binding to untransfected cells (gray), wild-type S-transfected cells (red), mutant S-transfected cells (light blue) are shown. **(C)** Titration of HCoV-HKU1 pseudoviruses with spike mutations corresponding to H501-008 binding residues as revealed by crystal structure in Fig. 2. Pseudovirus entry was quantified using RLU and plotted against log virus dilutions. All mutant pseudoviruses, except HCoV-HKU1-S-R447A (orange), were infectious, and thus their infectivity curves (multiple colors) overlap. **(D)** H501-008 (10 µg/mL) **n**eutralization assay of HCoV-HKU1 pseudoviruses with indicated spike mutations as revealed by crystal structure in Fig. 2. Data shown are mean ± s.d. of n = 2 independent experiments performed in technical triplicates. Statistical analysis was performed using one-way ANOVA; ****P <0.0001.

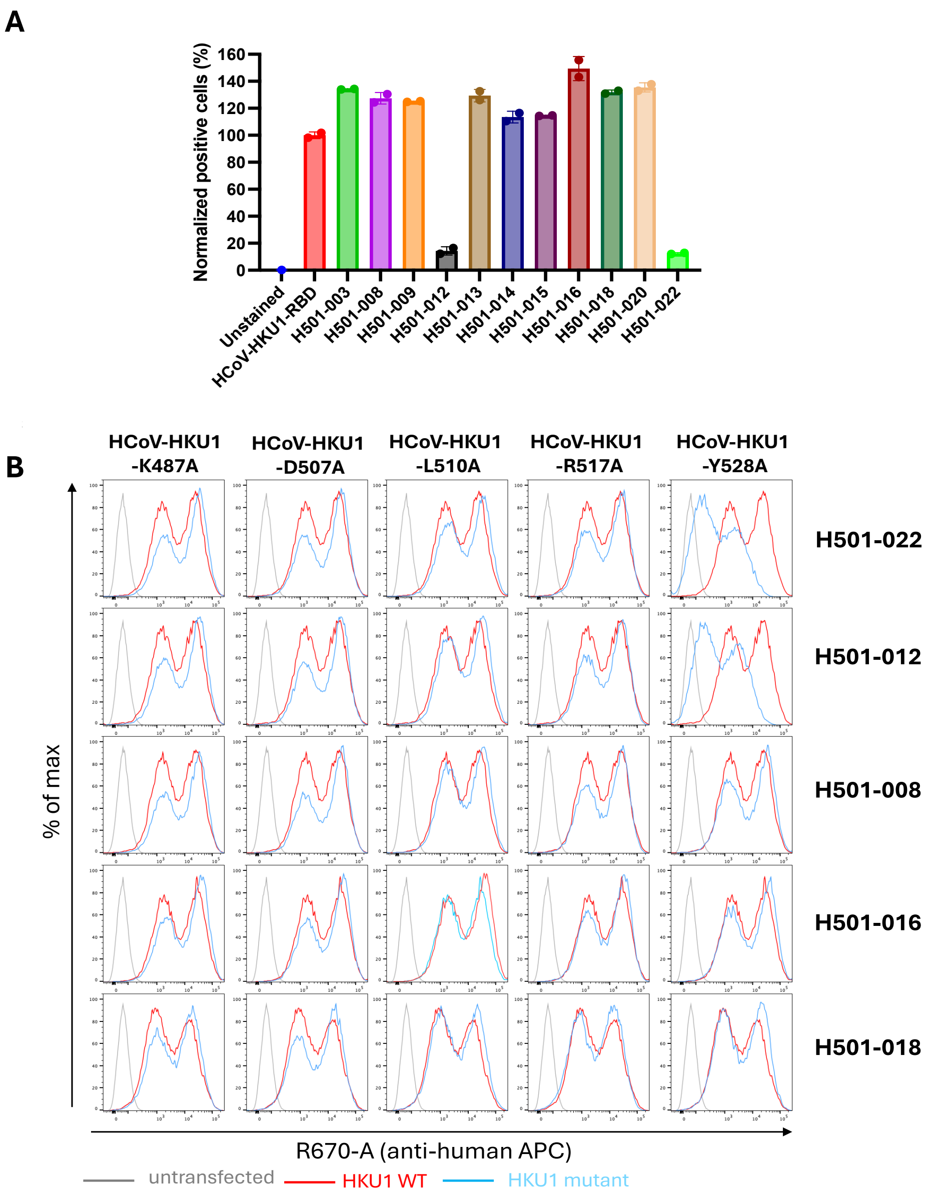


**Figure S9. HCoV-HKU1 RBD-specific mAbs H501-012 and -022 block S binding to TMPRSS2. (A)** HCoV-HKU1 RBD mAbs were analyzed for blocking S-TMPRSS2 interaction by flow cytometry analysis of cell surface-bound HCoV-HKU1 RBD-mAb complexes. HCoV-HKU1 RBD alone binding positive cells are set to 100% and the RBD-mAb complexes are normalized to RBD alone and represented as percent positive cells. **(B)** H501-012 and -022 were assessed for binding to cell surface-expressed HCoV HKU1 S with indicated TMPRSS2 binding site mutations, as revealed by crystal structure in Fig. 5. Percentage of maximum binding to untransfected cells (gray), wild-type S-transfected cells (red), mutant S-transfected cells (light blue) are shown.

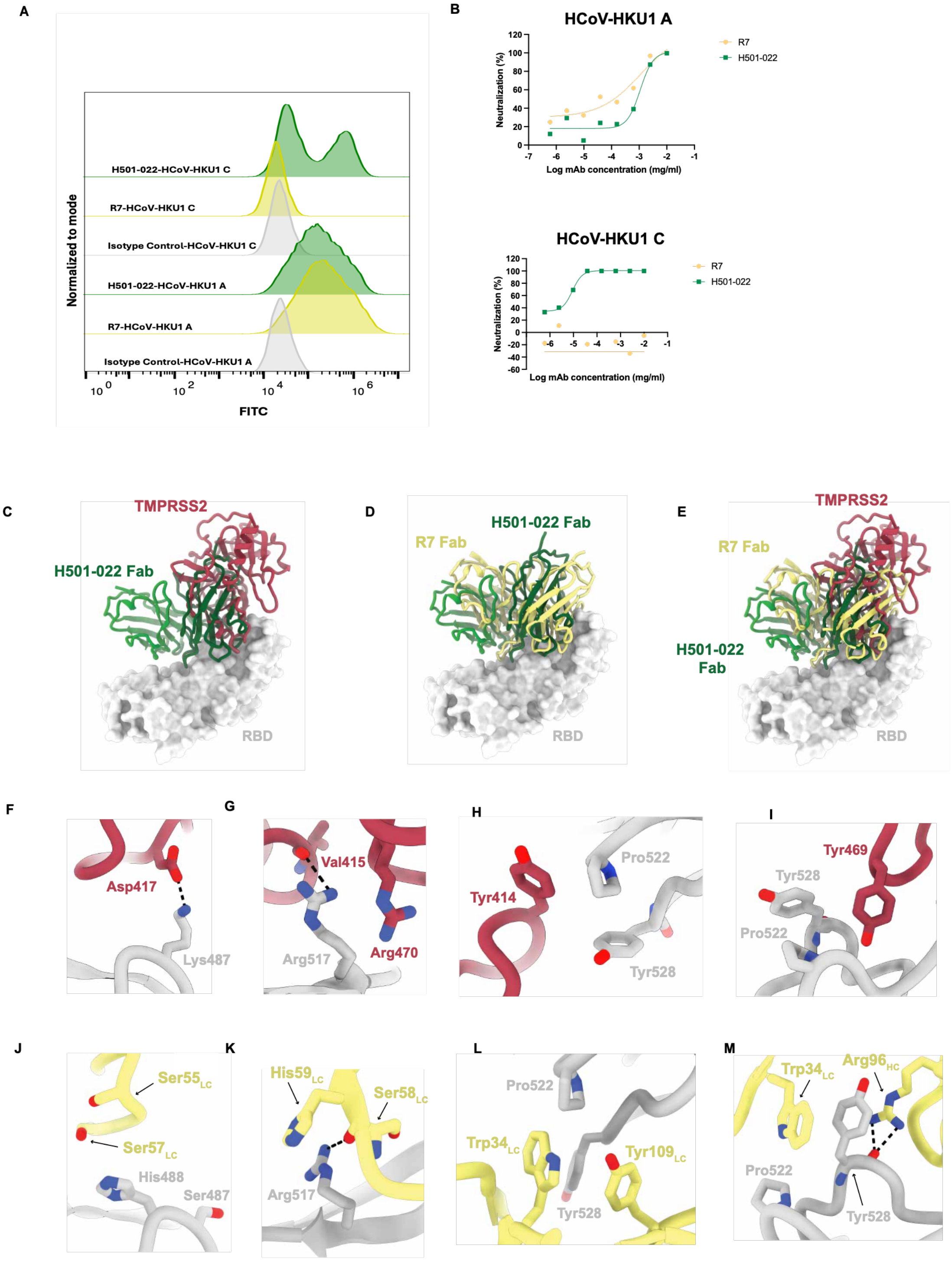


N

| | | Receptor Binding Domain | | | | | | | | | | | | | | | | | | | | | | | |
|---|---|---|---|---|---|---|---|---|---|---|---|---|---|---|---|---|---|---|---|---|---|---|---|---|---|
| Residue Number | | 465 | 487 | 488 | 501 | 508 | 509 | 510 | 515 | 517 | 518 | 521 | 522 | 525 | 526 | 527 | 528 | 529 | 530 | 531 | 532 | 536 | 537 | 538 | 539 |
| Amino Acid | HKU1 C S | H | K | S | Y | T | T | L | W | R | C | L | P | I | S | T | Y | S | P | N | T | K | K | V | V |
| | HKU1 A S | Y | S | H | Y | T | V | L | W | R | C | L | P | I | P | A | Y | D | P | R | S | K | K | S | L |
| BSA (Å$^2$) | H501-022 ($V_H + V_L$) | 39.1 | 0.0 | 0 | 0.0 | 0.0 | 52.3 | 73.0 | 42.8 | 62.4 | 3.0 | 19.5 | 51.7 | 33.0 | 44.4 | 20.1 | 129.2 | 22.4 | 80.6 | 74.9 | 8.3 | 50.9 | 90.8 | 17.4 | 2.8 |
| | TMPRSS2 | 0.0 | 25.6 | 31.1 | 0.0 | 4.5 | 64.1 | 111.3 | 55.7 | 87.3 | 12.4 | 60.7 | 25.9 | 0.0 | 0.0 | 0.6 | 153.1 | 43.3 | 14.4 | 43.5 | 19.5 | 0.0 | 0.0 | 0.0 | 0.0 |
| | R7 ($V_H + V_L$) | 37.1 | 0.0 | 8.6 | 0.0 | 53.8 | 46.7 | 126.8 | 46.2 | 76.2 | 0.0 | 25.4 | 50.1 | 10.5 | 52.6 | 10.2 | 137.5 | 19.2 | 7.1 | 138.9 | 0.0 | 0.0 | 49.5 | 0.0 | 14.9 |

**Figure S10. Comparison of RBD mAbs H501-022 and R7 HCoV-HKU1 S binding and neutralization.** **(A)** Cell surface HCoV-HKU1 spike (genotype A and C) binding assay using mAbs H501-022 and R7. Data represent mean fluorescent intensity (FITC) histograms. **(B)** H501-022 and R7 neutralization assay with HCoV-HKU1 pseudoviruses from genotypes A (upper panel) and C (lower panel). Neutralization curves were plotted as a nonlinear fit sigmoidal curve, normalized to cells and virus controls. (**C)** HCoV-HKU1 RBD (light gray) bound to H501-022 (green) from this study and TMPRSS2 (red, PDB 8Y8B). Structures were aligned to the RBD from this study. **(D)** HCoV-HKU1 RBD (light gray) bound to R7 (yellow, PDB 9JCY) and H501-022 (green). Structures were aligned to the RBD from this study. **(E)** Overlay of R7 (yellow, PDB 9JCY), TMPRSS2 (red, PDB 8Y8B), and H501-022 (green) bound to the RBD. Structures were aligned to the RBD from this study. **(F - M)** Residue interactions between RBD (gray) and TMPRSS2 (red) from PDB 8Y8B. **(J - M)** Residue interactions between RBD (gray) and R7 (yellow) from PDB 9JCY. **(N)** Table depicting sequence alignment between of HCoV-HKU1 A (GenBank Q5MQD0) and HCoV-HKU1 C (GenBank DQ339101) along with Buries Surface Area (BSA) analyses. BSA values are shown for HCoV-HKU1 C RBD bound to H501-022 (this study), HCoV-HKU1 C RBD bound to TMPRS22 (PDB 8Y8B) and HCoV-HKU1 A bound to R7 (PDB 9JCY). Values highlighted by darker colors indicate increased BSA contribution whereas values highlighted by lighter colors indicate less BSA contribution.

**HCoV-HKU1 RBD Footprint (H501-008, H501-018 and H501-022 contact Spike residues)**

```
                 441                                   480                                 520                                     564
Genotype C
ABD75513.1       PSSWNRRYGFGSFNLSSYDVVYSDHCFSVNSDFCPCADPSVVNSCVKSKPPSAICPAGTKYRHCDLD-TTLYVKNWCRCSCLPDPISTYSPNTCPQKKVVVGIGEHCPGLGINEEKCGTQLNHSS---
Q0ZME7.1         PSSWNRRYGFGSFNLSSYDVVYSDHCFSVNSDFCPCADPSVVNSCAKSKPPSAICPAGTKYRHCDLD-TTLYVKNWCRCSCLPDPISTYSPNTCPQKKVVVGIGEHCPGLGINEEKCGTQLNHSS---
ABD75505.1       PSSWNRRYGFGSFNLSSYDVVYSDHCFSVNSDFCPCADPSVVNSCVKSKPPSAICPAGTKYRHCDLD-TTLYVKNWCRCSCLPDPISTYSPNTCPQKKVVVGIGEHCPGLGINEEKCGTQLNHSS---
ABD75625.1       PSSWNRRYGFGSFNLSSYDVVYSDHCFSVNSDFCPCADPSVVNSCVKSKPPSAICPAGTKYRHCDLD-TTLYVKNWCRCSCLPDPISTYSPNTCPQKKVVVGIGEHCPGLGINEEKCGTQLNHSS---
AXT92529.1       PSSWNRRYGFGSFNVSSYDVVYSDHCFSVNSDFCPCADPSVVNSCVKSKPLSAICPAGTKYRHCDLD-TTLYVKNWCRCSCLPDPISTYSPNTCPQKKVVVGIGEHCPGLGINEEKCGTQLNHSS---
AXT92551.1       PSSWNRRYGFGSFNVSSYDVVYSDHCFSVNSDFCPCADPSVVNSCVKSKPLSAICPAGTKYRHCDLD-TTLYVKNWCRCSCLPDPISTYSPNTCPQKKVVVGIGEHCPGLGINEEKCGTQLNHSS---
URC25161.1       PSSWNRRYGFGSFNVSSYDVVYSDHCFSVNSDFCPCADPSVVNSCVKSKPLSAICPAGTKYRHCDLD-TTLYVKNWCRCSCLPDPISTYSPNTCPQKKVVVGIGEHCPGLGINEEKCGTQLNHSS---
AXT92528.1       PSSWNRRYGFGSFNVSSYDVVYSDHCFSVNSDFCPCADPSVVNSCVKSKPLSAICPAGTKYRHCDLD-TTLYVKNWCRCSCLPDPISTYSPNTCPQKKVVVGIGEHCPGLGINEEKCGTQLNHSS---
URC25188.1       PSSWNRRYGFGSFNVSSYDVVYSDHCFSVNSDFCPCADPSVVNSCVKSKPLSAICPAGTKYRHCDLD-TTLYVKNWCRCSCLPDPISTYSPNTCPQKKVVVGIGEHCPGLGINEEKCGTQLNHSS---
AGT17758.1       PSSWNRRYGFGSFNVSSYDVVYSDHCFSVNSDFCPCADPSVVNSCVKSKPLSAICPAGTKYRHCDVDTTTLYVNNWCRCSCLPDPISTYSPNTCPQKKVVVGIGEHCPGLGINEEKCGTQLNHSS--
WDE18934.1       PSSWNRRYGFSSFNVSSYDVVYSDHCFSVNSDFCPCADPSVVNSCVKSKPLSAICPAGTKYRHCDLDTTTLYVKNWCRCSCLPDPISTYSPNTCPQKKVVVGIGEHCPGLGINEEKCGTQLNHTS---
Genotype B
ABD75545.1       PSSWNRRYGFGSFNVSSYDVVYSDHCFSVNSDFCPCADRSVVNSCVKSKPPSAICPAGTKYRHCDLD-TTLYVKNWCRCSCLPDPISTYSPNTCPQKKVVVGIGEHCPGLGINEEKCGTQLNHSS---
ABD75617.1       PSSWNRRYGFGSFNVSSYDVVYSDHCFSVNSDFCPCADPSVVNSCVKSKPLSAICPAGTKYRHCDLD-TTLYVKNWCRCSCLPDPISTYSPNTCPQKKVVVGIGEHCPGLGINEEKCGTQLNHSS---
Q14EB0.1         PSSWNRRYGFGSFNVSSYDVVYSDHCFSVNSDFCPCADPSVVNSCVKSKPLSAICPAGTKYRHCDLD-TTLYVNNWCRCSCLPDPISTYSPNTCPQKKVVVGIGEHCPGLGINEEKCGTQLNHSS---
AXT92533.1       PSSWNRRYGFGSFNVSSYDVVYSDHCFSVNSDFCPCADPSVVNSCVKSKPLSAICPAGTKYRHCDLD-TTLYVKNWCRCSCLPDPISTYSPNTCPQKKVVVGIGEHCPGLGINEEKCGTQLNHSS---
AXT92559.1       PSSWNRRYGFGSFNVSSYDVVYSDHCFSVNSDFCPCADPSVVNSCVKSKPLSAICPAGTKYRHCDLD-TTLYVKNWCRCSCLPDPISTYSPNTCPQEKVVVGIGEHCPGLGINEEKCGTQLNHSS---
BBA20986.1       PSSWNRRYGFGSFNVSSYDVVYSDHCFSVNSDFCPCADPSVVNSCVKSKPLSAICPAGTKYRHCDLDTTTLYVKNWCRCSCLPDPISTYSPNTCPQKKVVVGIGEHCPGLGINEEKCGTQLNHTS---
AXT92548.1       PSSWNRRYGFGSFNVSSYDVVYSDHCFSVNSDFCPCADPSVVNSCVKSKPLSAICPAGTKYRHCDLDTTTLYVKNWCRCSCLPDPISTYSPNTCPQKKVVVGIGEHCPGLGINEEKCGTQLNHTS---
WDE18925.1       PSSWNRRYGFGSFNVSSYDVVYSDHCFSVNSDFCPCADPSVVNSCVKSKPLSAICPAGTKYRHCDLDTTTLYVKNWCRCSCLPDPISTYSPNTCPQKKVVVGIGEHCPGLGINEEKCGTQLNHTS---
AXT92540.1       PSSWNRRYGFGSFNVSSYDVVYSDHCFSVNSDFCPCADPSVVNSCVKSKPLSAICPAGTKYRHCDLDTTTLYVKNWCRCSCLPDPISTYSPNTCPQKKVVVGIGEHCPGLGINEEKCGTQLNHTS---
ARB07438.1       PSSWNRRYGFGSFNVSSYDVVYSDHCFSVNSDFCPCADPSVVNSCAKSKPLSAICPAGTKYRHCDVDSTTLYVNNWCRCSCLPDPISTYSPNTCPQKKVVVGIGEHCPGLGINEEKCGTQLNHSS---
WKE35756.1       PSSWNRRYGFGSFNVSSYDVVYSDHCFSVNSDFCPCADPSVVNSCDKSKPLSAICPTGTKYRHCDLD -TTLYVKNWCRCSCLPDPISTYSPNTCPQKKVVVGIGEHCPGLGINEEKCGTQLNHSS---
HKU1-B-This
study            PSSWNRRYGFGSFNVSSYDVVYSDHCFSVNSDFCPCADPSVVNSCDKSKPLSAICPTGTKYRHCDLD -TTLYVKNWCRCSCLPDPISTYSPNTCPQKKVVVGIGEHCPGLGINEEKCGTQLNHSS
Genotype A
YP_173238.1      PSSWNRRYGFNNFNLSSHSVVYSRYCFSVNNTFCPCAKPSFASSCKSHKPPSASCPIGTNYRSCEST  - TVLDHTDWCRCSCLPDPITAYDPRSCSQKKSLVGVGEHCAGFGVDEEKCGV-LDGSYNVS
AXT92550.1       PSSWNRRYGFNNFNLSSHSVVYSRYCFSVNNTFCPCAKPSFASSCKSHKPPSASCPIGTNYRSCEST  - TVLDRTDWCRCSCLPDPITAYDPRSCSQKKSLVGVGEHCAGFGVDEEKCGV-LDGSYNVS
ABD75601.1       PSSWNRRYGFNNFNLSSHSVVYSRYCFSVNNTFCPCAKPSFASSCKSHKPPSASCPIGTNYRSCEST  - TVLDHTDWCRCSCLPDPITAYDPRSCSQKKSLVGVGEHCAGFGVDEEKCGV-LDGSYNVS
WDE18950.1       PSSWNRRYGFNNFNLSSHSVVYSRYCFSVNNTFCPCANPSFASSCKSHKPPSASCPIGTNYRSCEST  - TVLDRTDWCRCSCLPDPITAYDPRSCSQKKSLVGVGEHCAGLGVDEEKCGV-LDGSYNAS
URC25143.1       PSSWNRRYGFNNFNLSSHSVVYSRYCFSVNNTFCPCANPSFASSCKSHKPPSASCPIGTNYRSCEST  - TVLDRTDWCRCSCLPDPITAYDPRSCSQKKSLVGVGEHCAGFGVDEEKCGV-LDGSYNAS
AMN88686.1       PSSWNRRYGFNNFNLSSHSVVYSRYCFSVNNTFCPCAKPSFASSCKSHKPPSASCPIGTNYRSCEST  -  TVLDHTDWCRCSCLPDPITAYDPRSCSQKKSLVGVGEHCAGFGVDEEKCGV-LDGSYNVS
AGT17777.1       PSSWNRRYGFNNFNLSSHSVVYSRYCFSVNNTFCPCAKPSFASSCKSHKPPSASCPIGTNYRSCEST  -  TVLDHTDWCRCSCLPDPITAYDPRSCSQKKSLVGVGEHCAGFGVDEEKCGV-LDGSYNVS
AXT92557.1       PSSWNRRYGFNNFNLSSHSVVYSRYCFSVNNTFCPCAKPSFASSCKSHKPPSASCPIGTNYRSCEST  -  TVLDHTDWCRCSCLPDPITAYDPRSCSQKKSLVGVGEHCAGFGVDEEKCGV-LDGSYNVS
AGW27872.1       PSSWNRRYGFNNFNLSSHSVVYSRYCFSVNNTFCPCAKPSFASSCKSHKPPSASCPIGTNYRSCEST  -  TVLDRTDWCRCSCLPDPITAYDPRSCSQKKSLVGVGEHCAGFGVDEEKCGV-LDGSYNAS
URC25127.1       PSSWNRRYGFNNFNLSSHSVVYSRYCFSVNNTFCPCANPSFASSCKSHKPPSASCPIGTNYRSCEST -  TVLDRTDWCRCSCLPDPITAYDPRSCSQKKSLVGVGEHCAGFGVDEEKCGV-LDGSYNAS
AMN88694.1       PSSWNRRYGFNNFNLSSHSVVYSRYCFSVNNTFCPCAKPSFASSCKSHKPPSASCPIGTNYRSCEST -   TVLDHTDWCRCSCLPDPITAYDPRSCSQKKSLVGVGEHCAGFGVDEEKCGV-LDGSYNVS
```

**Figure S11. mAbs H501-008, -018 and -022 contact residues across HCoV-HKU1 strains.** HCoV-HKU1 genotypes A, B, and C RBD alignment highlighting H501-008, H501-018, and H501-022 contact residues in red, purple, and green, respectively. HCoV-HKU1 S sequences were retrieved from the NCBI, GenBank. In MEGA11, these sequences were aligned using the MUSCLE algorithm together with the sequence of the HCoV-HKU1 live virus used in this study.

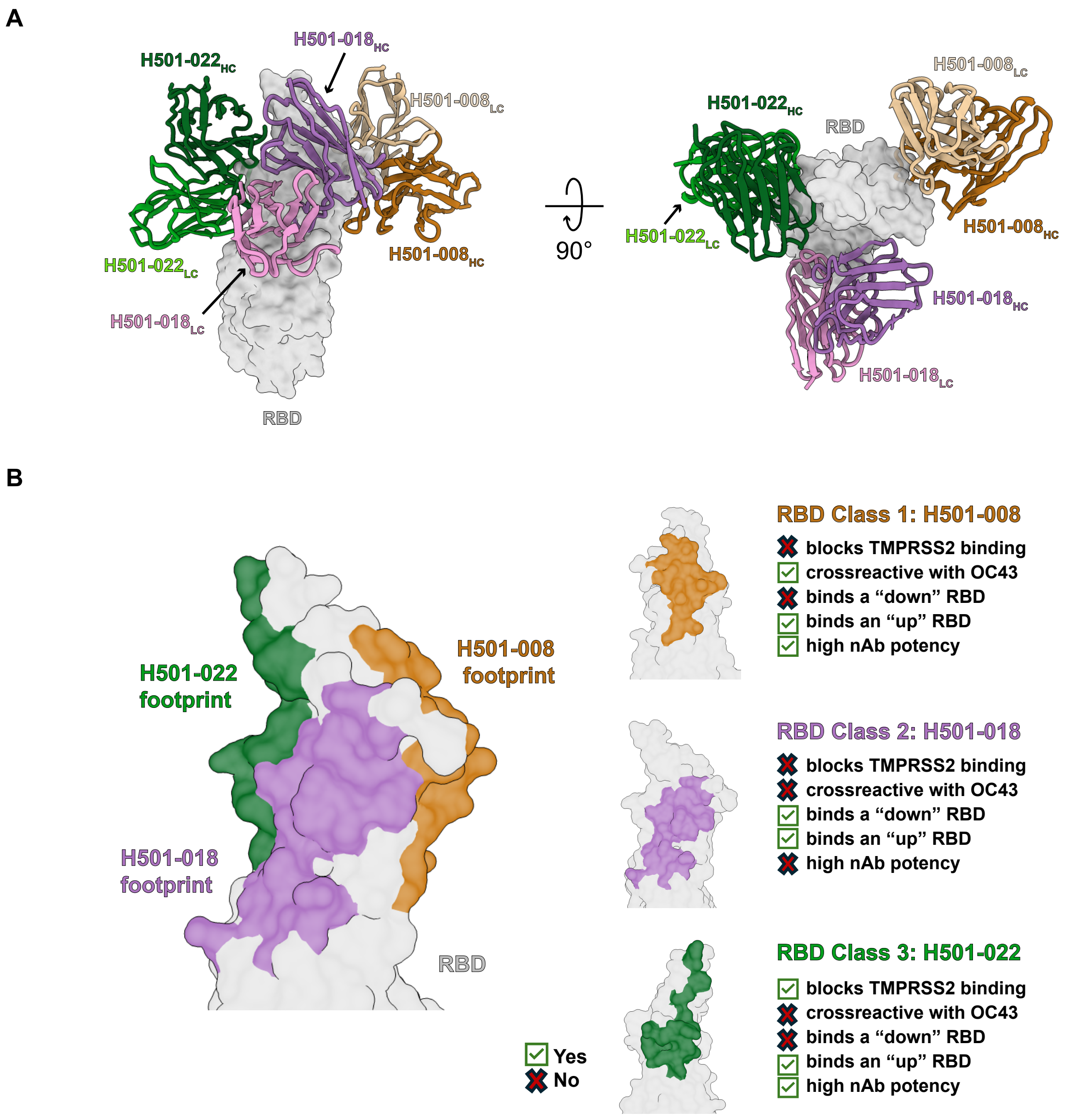


**Figure S12. Comparison of mAbs H501-008, -018, and -022 bound to HCoV-HKU1 RBD. (A)** Heavy chains and light chains from H501-008, -018, and -022 are aligned based on the RBD. RBD is shown as a surface representation, and Fabs are shown as ribbon representations. **(B)** Summarized characteristics of H501-008, H501-018, and H501-022. RBD epitopes are color-coded based on their respective antibodies. Colors are as follows: RBD (light gray), H501-008 $V_H$ (orange), H501-008 $V_L$ (tan), H501-008 epitope (orange), H501-018 $V_H$ (purple), H501-018 $V_L$ (pink), H501-018 epitope (purple), H501-022 $V_H$ (green), H501-022 $V_L$ (light green), H501-022 epitope (green).

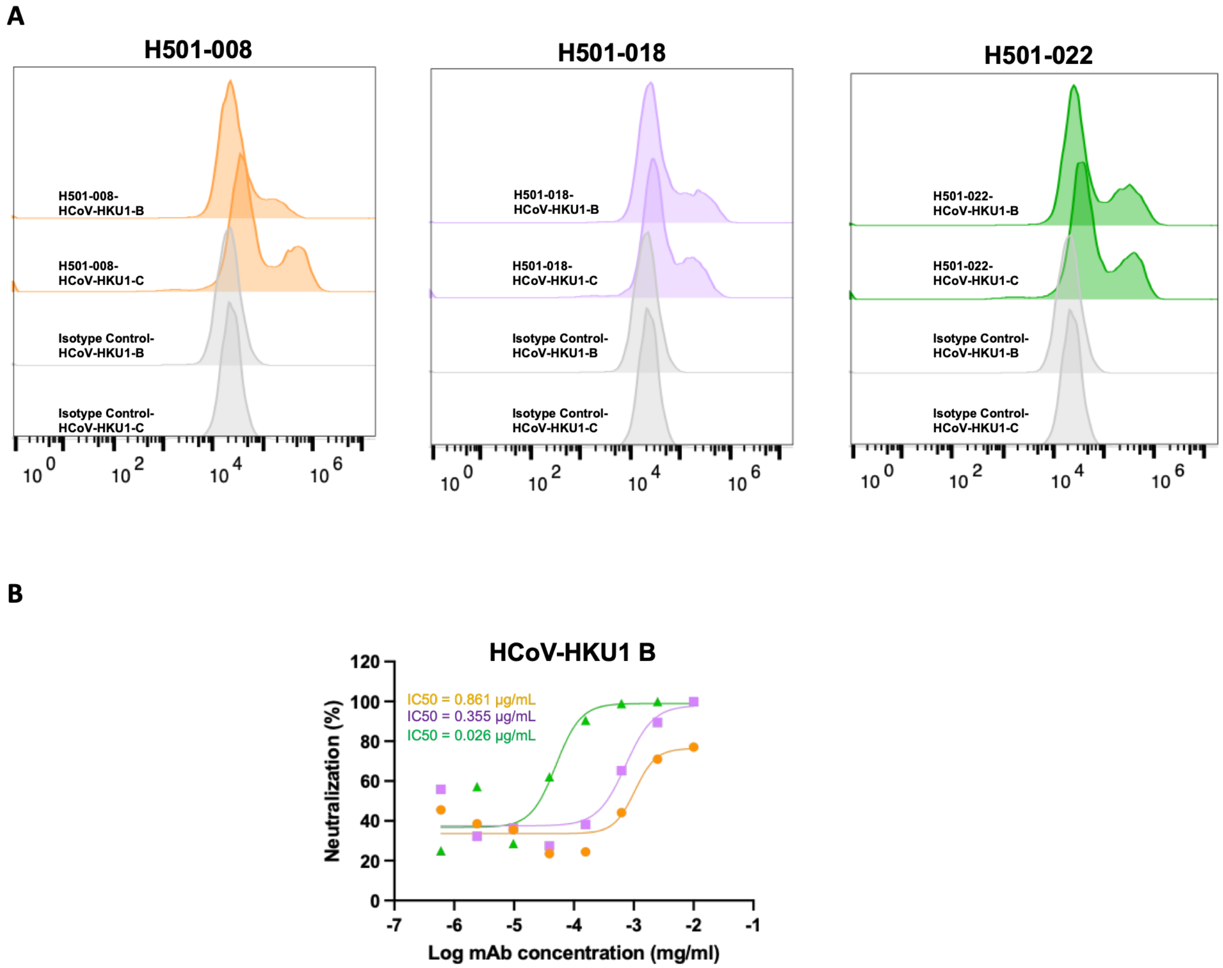


**Figure S13. HCoV-HKU1-B vs. HCoV-HKU1-C: Comparison of cell surface mAb binding and neutralization. (A)** Cell surface mAb binding assay of HCoV-HKU1 S (genotypes B and C) using HCoV-HKU1 mAbs H501-008 (orange), H501-018 (purple), and H501-022 (green). Data represent mean fluorescent intensity histograms. **(B)** HCoV-HKU1-B pseudovirus neutralization assay with HCoV-HKU1 mAbs H501-008, H501-018 and H501-022. Neutralization curves were plotted as a nonlinear fit sigmoidal curve, normalized to cells and virus controls.

**A**

**B**

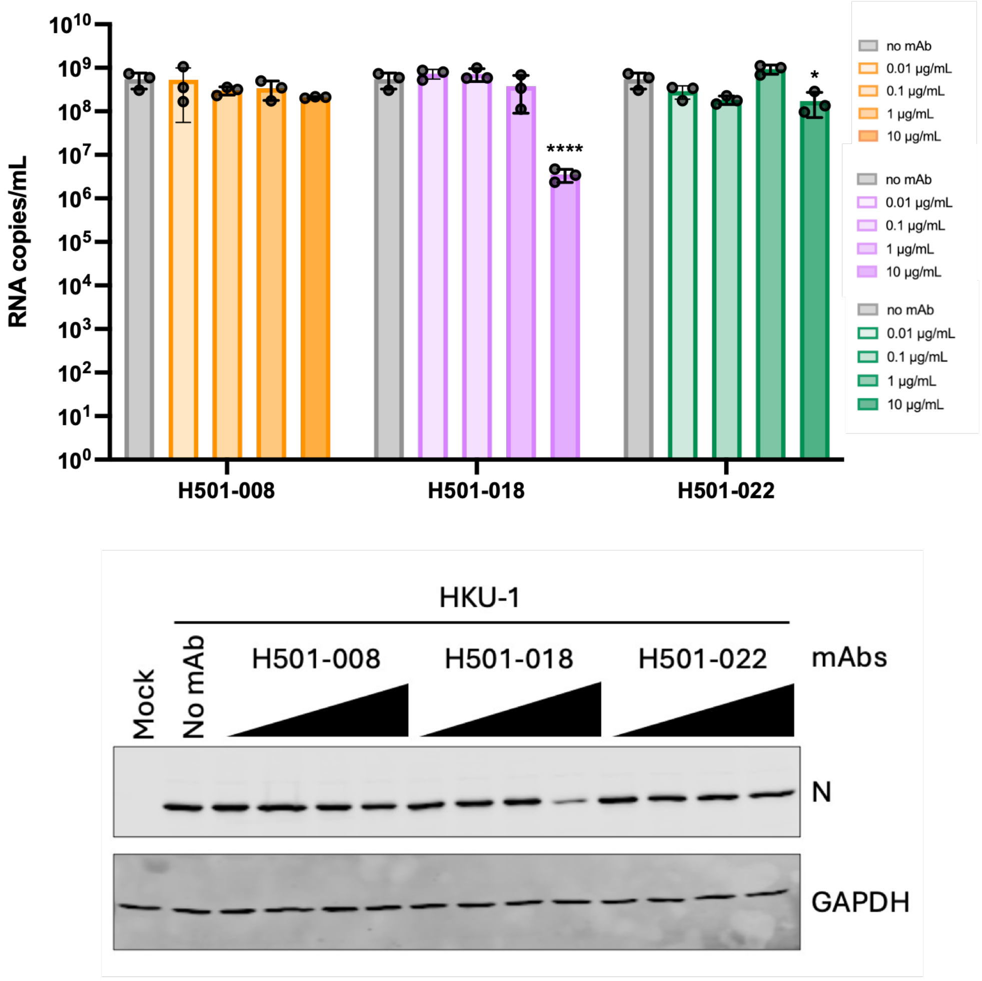


**Figure S14. HCoV-HKU1 live virus neutralization assay in NHBE cells. (A)** Concentration-dependent neutralization effect H501-008 (orange), -018 (purple), and -022 (green) against HCoV-HKU1 live virus infection in NHBE cells (Donor 2) was analyzed by RT-PCR of viral nucleocapsid (N) gene. Twenty-four hours post-infection, RNA copies/ ml was calculated with reference to the standard curve generated using in vitro–transcribed RNA corresponding to the HKU1 N gene. Error bars denote mean ± SD. RNA copies at Individual time points were compared to no mAb control using one-way ANOVA with Dunnett's multiple comparison. *P ≤ 0.05, ****P ≤ 0.0001. **(B)** HCoV-HKU1 was pre-incubated with increasing concentrations of mAbs prior to infection, and viral N protein expression was assessed by Western blotting of cell lysates collected at 96 hours post-infection. GAPDH was used as an internal control.
